# PML mutants resistant to arsenic induced degradation fail to generate the appropriate SUMO and ubiquitin signals required for RNF4 and p97 recruitment

**DOI:** 10.1101/2023.01.15.524136

**Authors:** Ellis G. Jaffray, Michael H. Tatham, Alejandro Rojas-Fernandez, Adel Ibrahim, Graeme Ball, Ronald T. Hay

## Abstract

Arsenic is an effective treatment for Acute Promyelocytic Leukaemia as it induces degradation of the Promyelocytic Leukaemia (PML) – retinoic acid receptor alpha (RARA) oncogenic fusion protein. Some patients relapse with arsenic resistant disease because of missense mutations in PML. To determine the mechanistic basis for arsenic resistance we reconstituted PML-/- cells with YFP fusions of wild type (WT) and two mutant forms of PMLV found in patients refractory to arsenic, A216T and L217F. Both mutants formed PML bodies that were larger, but less numerous than WT and neither responded to arsenic by degradation. Analysis of immunoprecipitated PML bodies indicated that while WT PML experiences increased SUMO1, SUMO2/3 and ubiquitin conjugation, A216T PML is almost completely unresponsive and therefore does not recruit the SUMO targeted ubiquitin E3 ligase RNF4. Compared to WT PML, L217F PML was modified to a similar extent by SUMO2 but not SUMO1 and although it recruited RNF4, it failed to develop the appropriate poly-ubiquitin conjugates required to recruit the segregase p97, which is essential for PML degradation.

## INTRODUCTION

Acute Promyelocytic Leukaemia (APL) is caused by a reciprocal chromosomal translocation t(15;17) that fuses the genes encoding the Promyelocytic Leukaemia (PML) protein and the retinoic acid receptor alpha (RARA) (de The, Lavau et al. 1991, Kakizuka, Miller et al. 1991). This generates the PML-RARA oncoprotein that deregulates transcriptional programmes required for the differentiation of haematopoetic progenitor cells. Therefore, differentiation is blocked and promyelocytes accumulate causing leukaemia (Grignani, Ferrucci et al. 1993, Kwok, Zeisig et al. 2006, Martens, Brinkman et al. 2010, Mikesch, Gronemeyer et al. 2010, Tan, Wang et al. 2021). In healthy cells the PML gene is alternatively spliced to generate multiple mRNAs that encode seven protein isoforms (I-VII) that differ in their C-terminal regions.

PML is a member of the Tripartite Motif (TRIM) family of proteins and is also known as TRIM19. Located in the N-terminal region of PML and present in all PML isoforms and PML-RARA fusions, the highly conserved TRIM is composed of a RING domain, two zinc co-ordinating B-boxes and a coiled-coil that mediates dimerization (Bernardi and Pandolfi 2007). In cells with wild type (WT) PML genes, the protein is associated with non-membranous nuclear structures known as PML nuclear bodies, that also accumulate a variety of other proteins including Small Ubiquitin-like Modifiers (SUMOs), p53, Daxx and SP100. In cells containing a PML-RARA fusion protein, PML and PML-RARA are co-associated in a large number of tiny nuclear speckles (Dyck, Maul et al. 1994, Weis, Rambaud et al. 1994). In the past, treatment options were limited and APL was a disease with a very poor prognosis. However, the majority of patients are now cured by a combination therapy consisting of arsenic trioxide (referred to herein as As) and *all trans* retinoic acid (ATRA) (Wang and Chen 2008, Lo-Coco, Avvisati et al. 2013, Mi, Chen et al. 2015). These induce degradation of the PML-RARA oncogene and allows the WT version of RARA to initiate a transcriptional programme that drives the accumulated promyelocytes down a pathway of terminal differentiation that ultimately cures the disease. Arsenic alone induces degradation of both the unfused PML and PML-RARA, identifying PML as the target of As. Treatment with As leads to rapid multisite modification of PML and PML-RARA with SUMO (Muller, Matunis et al. 1998). SUMO modified PML and PML-RARA then serves to recruit the SUMO Targeted Ubiquitin E3 ligase (STUbL) RING Finger Protein 4 (RNF4) (Perry, Tainer et al. 2008, Geoffroy and Hay 2009). Recruitment of RNF4 to SUMO modified PML and PML-RARA present in PML nuclear bodies is mediated by multiple SUMO Interaction Motifs (SIMs) present in the N-terminal region of the protein. The high local concentration of RNF4 leads to dimerization of the C-terminal RING domains and activation of the ubiquitin E3 ligase activity of the protein (Rojas-Fernandez, Plechanovova et al. 2014). RNF4 mediated ubiquitination of SUMO modified PML and PML-RARA ultimately leads to their proteolytic degradation by the proteasome (Lallemand-Breitenbach, Jeanne et al. 2008, Tatham, Geoffroy et al. 2008). Recently, an additional step required for arsenic induced degradation of PML and PML-RARA has been uncovered that involves the p97/VCP segregase (Jaffray, Tatham et al. 2023). RNF4 mediated ubiquitination of PML leads to recruitment of UFD1-NPLOC4-p97 which extracts ubiquitinated PML and presumably feeds the modified protein to the proteasome where it is proteolytically degraded.

While the majority of APL patients treated with As are cured, a small proportion of these patients relapse and present with arsenic resistant disease. In some cases resistance may be a consequence of metabolic reprogramming linked to expression of additional known oncogenes (Madan, Shyamsunder et al. 2016, Iaccarino, Ottone et al. 2019), but in many patients, arsenic resistance is a result of missense mutations in PML clustered between residues L211 and S220 in B-box 2. These mutations are not only found in the PML-RARA oncogene (Goto, Tomita et al. 2011, Chendamarai, Ganesan et al. 2015, Lou, Ma et al. 2015, Iaccarino, Ottone et al. 2016, Liu, Zhu et al. 2016) but also in the unfused version of PML (Lehmann-Che,= Bally et al. 2014, Zhu, Qin et al. 2014, Iaccarino, Ottone et al. 2016). These mutations are undetectable at initial diagnosis, suggesting that cells expressing these mutations are selected for by arsenic treatment (Alfonso, Iaccarino et al. 2019, Balasundaram, Ganesan et al. 2022).

To establish the mechanistic basis for the arsenic resistance we investigated the properties of two mutations in PML that responded differently to As. While both mutations were resistant to As-induced degradation, the A216T mutant was severely compromised for SUMO modification, failed to recruit RNF4 and as a result was not ubiquitinated. In contrast, like WT PML the L217F mutant was modified by SUMO2/3, recruited RNF4, and was ubiquitinated in response to As. However, SUMO1 modification was compromised and the ubiquitin signal was not sufficient to recruit the p97 segregase that is required to extract PML from nuclear bodies. Thus, while two mutations that are adjacent in the PML B-box both inhibit As-induced PML degradation, mechanistic failure is at different steps of the degradation pathway.

## RESULTS

### PML mutants A216T and L217F form atypical PML bodies in U2OS cells

Mutations in PML and PML-RARA that arise in APL patients with arsenic resistant disease are a unique resource as these mutations are selected in vivo to be resistant to arsenic while still retaining the biological functions of PML that contribute to the disease phenotype. The majority of these mutations are located to a 10 amino acid region (L211-S220) in B-box 2 of PML that encompasses, but never includes, the zinc co-ordinating residues C212 and C215 (Fig. 1A). As PML is the direct target of arsenic and mutations from patients with arsenic resistant disease can also be detected in the non-rearranged allele of PML (Lehmann-Che, Bally et al. 2014, Iaccarino, Ottone et al. 2016), we chose to reconstitute U2OS PML-/- cells with wild-type (WT) or mutant versions of PMLV N-terminally fused to YFP (Jaffray, Tatham et al. 2023). In testing a number of PML variants, we found that they fell into two main groups exemplified by the most frequently identified mutations in patients; A216T and L217F. U2OS cells expressing YFP fused forms of WT, A216T and L217F PMLV were isolated by FACS and the derived cell lines analysed by high content microscopy. Images of these cells show that while the total YFP fluorescence signal was similar for WT, A216T and L217F, the average size and number of PML bodies per cell varied (Fig. 1B). Quantitative analysis of thousands of cells showed that the average expression levels, measured by YFP-PML intensity per cell, were similar for all three PML forms (Fig. 1C). However, cells expressing A216T and L217F PMLV had on average approximately half the number of PML bodies per cell (median = 4) compared to WT (median = 8) (Fig. 1C). This was contrasted by both mutants showing larger than WT PML bodies according to their measured cross-sectional area (Fig. 1C). This ∼20-25% larger area equates to almost twice the volume when considering a spherical PML body. These results show that A216T and L217F mutations to PML reduce PML body number and increase PML body size in untreated cells.

**Figure 1.**
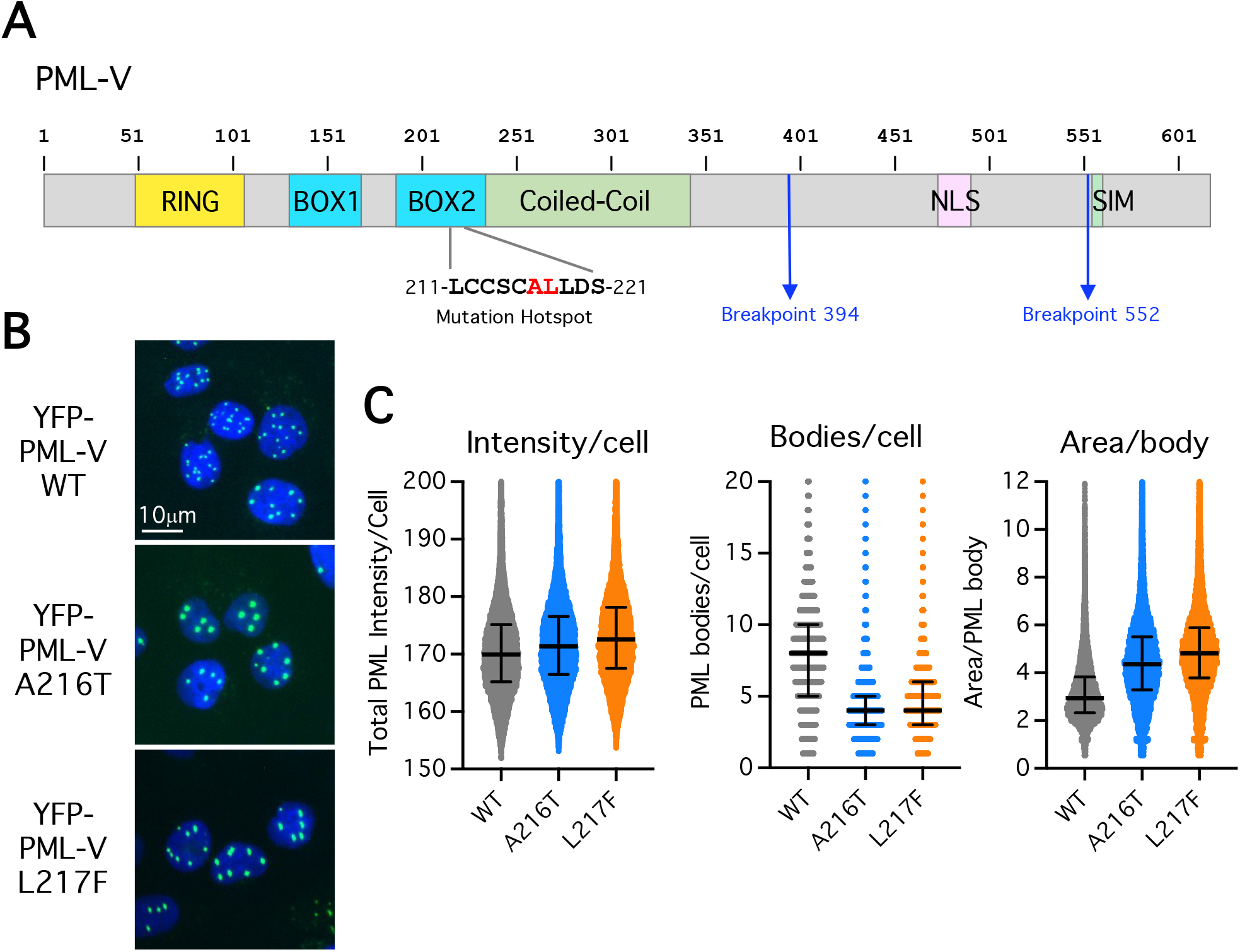
Characterization of PML bodies in U2OS PML-/- cell lines expressing YFP-fusions of WT, A216T and L217F forms of PML-V. A. Schematic depiction of PML-V primary structure indicating the major domains, regions of common mutations associated with arsenic treatment resistance, and the common breakpoints leading to PML-RARA fusions. B. Representative images of the indicated PML-/- U2OS + YFP-PML-V cell lines C. High content imaging data summarizing the calculated values for total PML intensity per cell, PML body number per cell and area per PML body for each cell line. Scatter plots are shown with median (solid line) and quartiles (error bars). n=38411 (wt), 35210 (A216T) and 41288 (L217F).

### PML mutants A216T and L217F are not degraded in response to arsenic in U2OS cells

Arsenic exposure is responsible for dramatic changes to the number, morphology and content of PML bodies (Geoffroy et al., 2010; Jeanne et al., 2010). To determine how these parameters were influenced by mutations in PML, the cells expressing WT, A216T or L217F YFP-PMLV were either untreated or treated with 1μM arsenic and monitored by time-lapse fluorescence microscopy over a 15 hr period. At least 9 movies were collected for each condition and an automated, quantitative pipeline (Jaffray, Tatham et al. 2023) was used to follow PML body size, number and total YFP-PMLV fluorescence intensity over time (see Supplementary movies 1-6 for representative time-course data). Representative images from the cells at 0 and 15 hr (Fig. 2A) showed the expected reduction in the number and total intensity of nuclear bodies in WT PMLV expressing cells, but not in cells expressing A216T or L217F PMLV (Fig. 2A). Averaged values of these metrics across all time courses indicated that PML body size did not significantly change over time after arsenic treatment for any of the PML forms studied (Fig. 2B). Furthermore, the number of bodies per cell and normalized YFP-PML intensity was reduced during arsenic exposure for WT PML but not for either mutant (Fig. 2B). More specifically, after 15 hr As there was a >4 fold reduction in the number of PML bodies per cell and >3 fold reduction in YFP-PMLV intensity for the WT PMLV, while little or no change was observed for A216T and L217F mutants (Fig. 2C). Western-blot analysis of extracts gathered from cells exposed to arsenic over a 24 hr time-course showed that as expected, WT PML rapidly shifts to a higher molecular weight species and is completely degraded by 24 hr (Fig. 2D). In contrast, A216T PML does not appear to shift to a higher molecular weight form and its levels do not change over the 24 hr period. Unlike A216T, L217F PML does shift to higher molecular weight species, but it is not degraded in response to arsenic after 24 hr (Fig. 2D).

**Figure 2.**
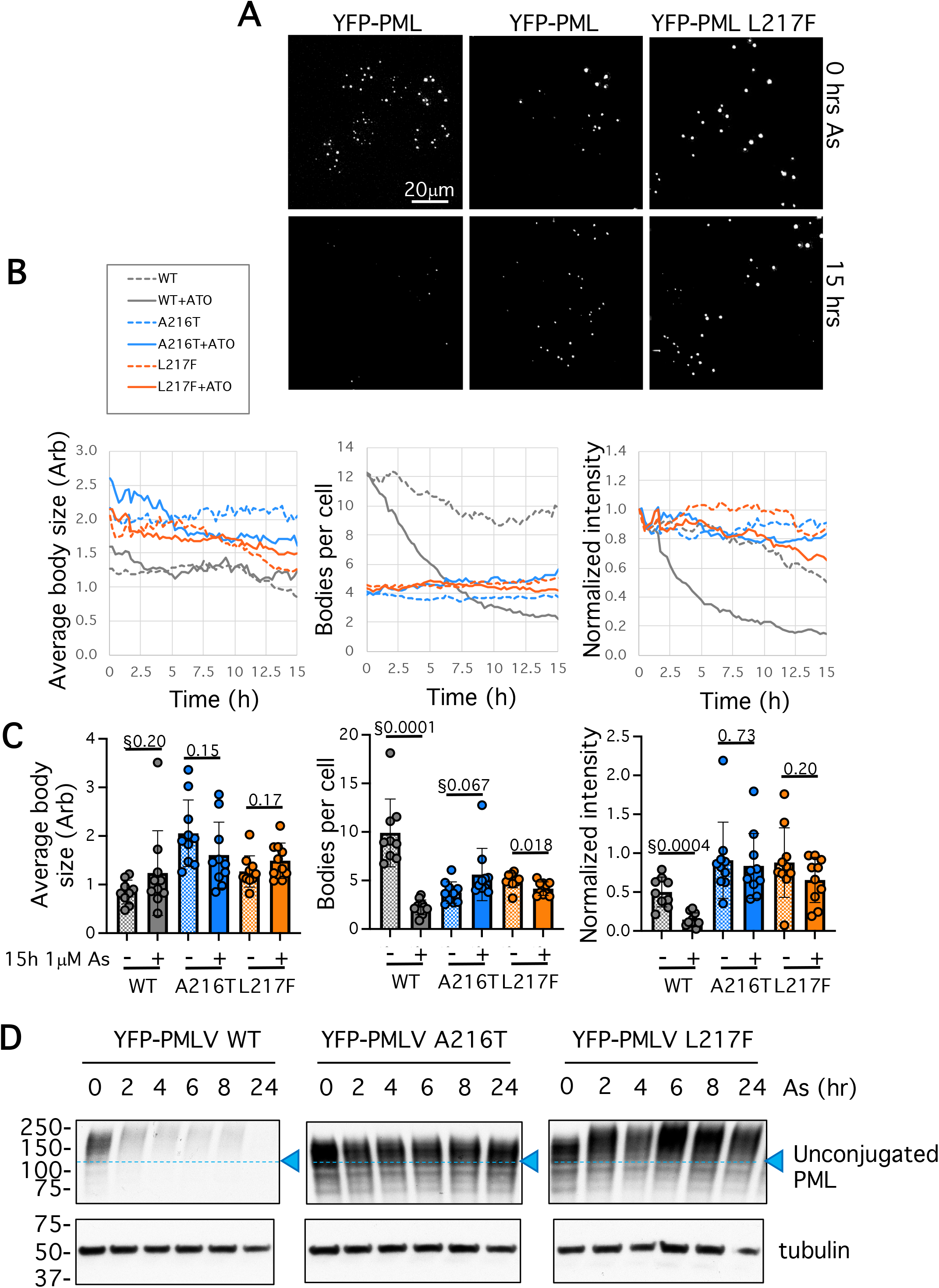
A216T and L217F mutants of PML are not rapidly degraded in response to arsenic treatment in vitro. A. YFP fluorescence image stills from one live cell analysis of each YFP-PML cell type at both 0h and 15h treatment with 1μM arsenic. B. Averaged data of 9 or 10 fields of view (median 12 cells per field) followed over a 15 hour time course of treatment or not with 1μM arsenic (15 minute intervals). Intensity measurements are normalized by the t=0hr values. C. Summary statistics (n=9 or 10) for 15hr data comparing 1μM arsenic treated cells with untreated cells. P-values are unpaired, two-tailed student’s T-tests using Welch’s correction for unequal variances where appropriate (§). See Supplementary movie files 1-6 for real-time data of a single representative field. D. Anti-PML Western-blots of fractionated whole cell extracts showing time-courses of the response of YFP- PML-V variants to 1μM arsenic exposure over 24 hours.

### In response to arsenic, WT and L217F PML recruit RNF4 but the A216T mutant does not

Although A216T does not appear to respond to arsenic by altered post-translational modification, both the WT and L217F forms display reduced electrophoretic mobility upon exposure of cells to arsenic (Fig. 2D). To better understand the nature of the modification status of all PML variants, nuclear bodies were purified from WT, A216T and L217F YFP-PMLV expressing cells (Jaffray, Tatham et al. 2023) both before and after treatment with 1μM As for 2h and analysed by Western blot for PML, SUMO1 or SUMO2/3. Consistent with the analysis of PML from whole cell extracts (Fig. 2D), purified YFP-PMLV WT and L217F shifted to higher molecular weight species in response to arsenic treatment for 2 hrs, whereas this response was almost completely blunted for A216T (Fig. 3A). SUMO1 modification of all three PML forms in untreated cells appears to be similar, but in response to arsenic, WT YFP-PMLV undergoes a much greater increase in SUMO1 modification than either A216T or L217F mutants. As with SUMO1, in untreated cells conjugation of all three PML forms to SUMO2/3 is broadly comparable (Fig. 3A). However, there are some notable differences in the SUMO2/3 response to arsenic in comparison with SUMO1. Firstly, the scale of the As-induced increase in SUMO2/3 conjugation to WT-PML is not as large as the relative increase in SUMO1. Secondly, both mutants show a SUMO2/3 conjugation response to As treatment, albeit weakest for the A216T mutant (Fig. 3A).

**Figure 3.**
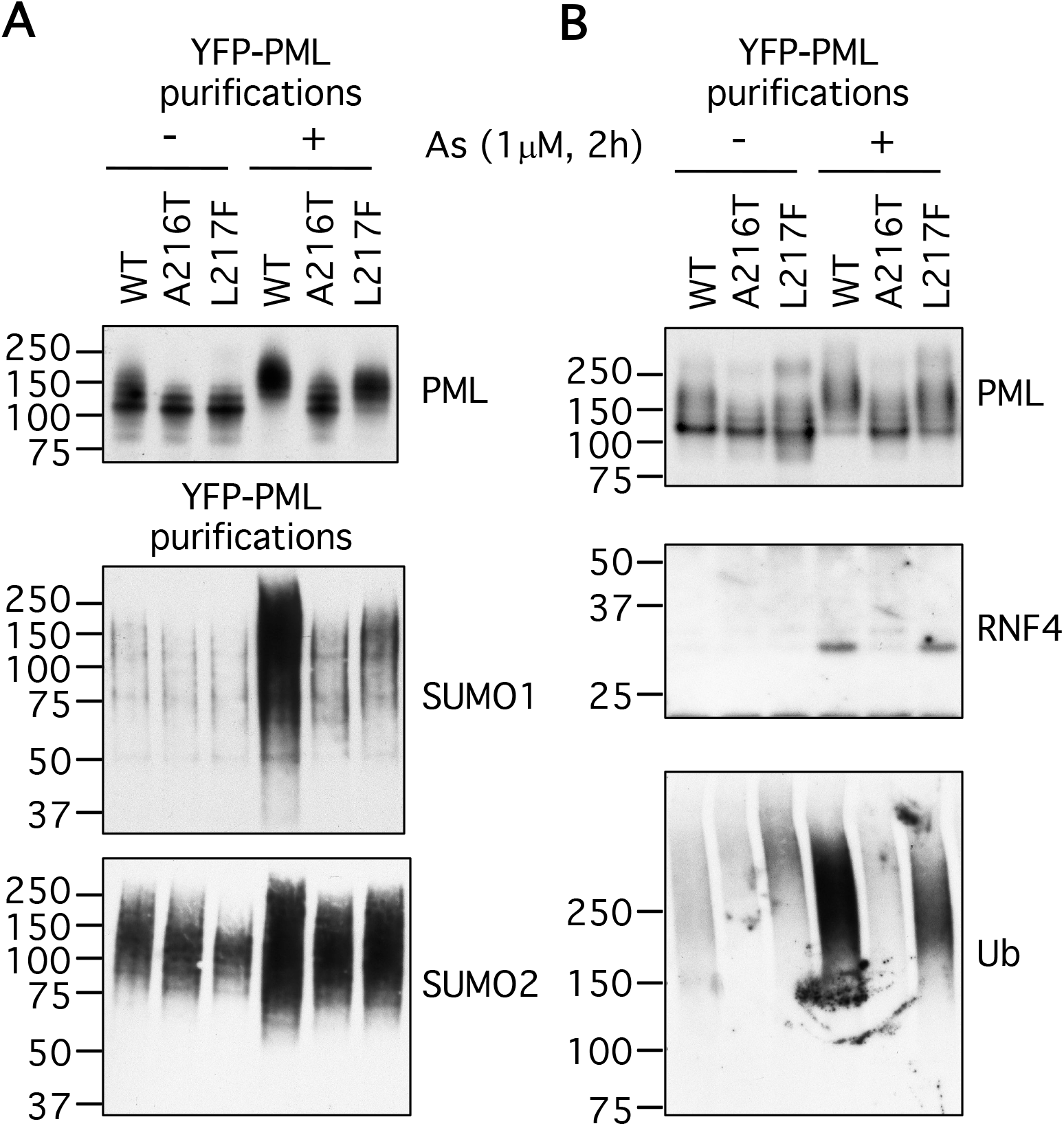
In response to arsenic A216T PML fails to recruit RNF4. Two separate purifications of YFP-PMLV WT, A216T and L217F (A+B): PML bodies were enriched from U2OS PML-/- + YFP-PMLV WT, A216T or L217F cells, either nontreated (NT) or treated with 1μM arsenic (1μM As 2h) for 2 hr. Bound material was eluted and analysed by Western blotting using antibodies to (A) PML, SUMO1, SUMO2/3 and (B) PML, RNF4 and ubiquitin.

Enhanced SUMO modification of PML is responsible for the recruitment of the STUbL RNF4 that mediates PML ubiquitination prior to degradation by the proteasome. To study RNF4 recruitment in response to arsenic, nuclear bodies from WT, A216T and L217F YFP-PMLV expressing cells were again purified and the associated RNF4 determined by Western blotting (Fig. 3B). RNF4 was not stably associated with nuclear bodies in the absence of arsenic, but was detected in purifications containing WT and L217F YFP-PMLV forms after arsenic treatment (Fig. 3B). By this method RNF4 was not detectable in A216T YFP-PMLV purifications in response to arsenic (Fig. 3B). The YFP-PMLV purifications were then analysed for ubiquitin modification by Western blotting with an antibody that detects ubiquitin chains. This revealed that WT YFP-PMLV was extensively ubiquitinated in response to arsenic and, consistent with the failure to recruit RNF4, increase in the ubiquitination of A216T YFP-PMLV was not detected (Fig. 3B). Interestingly, although L217F YFP-PMLV recruited RNF4, the increase in ubiquitin chains was not apparently as great as for the WT protein, and the modified species had a lower molecular weight (Fig. 3B).

The arsenic-induced recruitment of SUMO1, SUMO2/3 and RNF4 to PML bodies was also analysed by fluorescence microscopy for the three different cell lines. This confirmed that colocalization of SUMO1, SUMO2/3 and RNF4 with PML was increased for the WT protein in response to arsenic (Supplementary Figs. 1-3). Consistent with the Western blot data (Fig. 3A+B), the A216T PML mutant showed little response to arsenic in terms of recruitment of SUMO1, SUMO2/3 or RNF4. The L217F PML variant showed lower than WT recruitment of SUMO1 and SUMO2/3 in response to arsenic (Supplementary Fig. 1&2), although RNF4 levels appeared similar (Supplementary Fig. 3). In this experiment, RNF4 appears to colocalise with PML nuclear bodies prior to arsenic treatment, but this increases in response to arsenic for both WT and L217F and less so for A216T (Supplementary Fig. 3). Thus, both Western blotting and fluorescence microscopy indicate that the inability of A216T PML to be degraded is the consequence of insufficient SUMO conjugation which fails to recruit RNF4, and therefore is not ubiquitinated. This was not the case for L217F which could recruit RNF4 and induce ubiquitination, but was still resistant to degradation.

### In response to arsenic L217F PML fails to recruit the p97 segregase

Recently we determined that the p97 segregase was required to extract ubiquitinated PML from nuclear bodies prior to degradation by the proteasome (Jaffray, Tatham et al. 2023). To determine if the mutants were compromised for p97 extraction we analysed WT, A216T and L217F YFP-PMLV expressing cells by fluorescence microscopy 0, 2, 4 and 6 hr after arsenic treatment. p97 recruitment was determined by evaluating the extent of co-localisation between antibody stained p97 and PML marked by YFP fluorescence. As p97 is a highly abundant protein we employed a pre-extraction procedure to release the bulk of the soluble p97 from the cells before fixation. In WT YFP-PMLV untreated cells p97 was associated with only a small proportion of PML bodies, whereas after 6 hr arsenic treatment a larger fraction of PML bodies had associated p97 (Supplementary Fig. 4). In cells expressing A216T and L217F YFP-PMLV, arsenic treatment did not appear to increase the association of p97 with PML bodies (Fig. 4A, Supplementary Fig. 4). To quantify this, the percentage of PML bodies co-localised with p97 in WT, A216T and L217F YFP- PMLV expressing cells was determined at 0, 2, 4 and 6 hr after arsenic treatment (Fig. 4B). In untreated WT YFP-PMLV expressing cells about 20% of PML bodies were associated with p97, which increased to over 60% at 2, 4 and 6 hr after arsenic treatment. In untreated cells expressing A216T and L217F YFP-PMLV, p97 association was less than 10% and this did not increase in response to arsenic (Fig. 4B). Statistical analysis of the 0 hr and 6 hr time-points data revealed that there was a substantial and statistically significant recruitment of p97 to PML bodies in cells expressing WT YFP-PMLV, but not in cells expressing A216T or L217F YFP-PMLV (Fig. 4C). Changes in the expression levels of p97 in the different cell lines, or upon arsenic treatment could not explain these differences (Supplementary Fig. 5). Thus, although L217F YFP-PMLV recruits SUMO and RNF4 to generate ubiquitin conjugates, this signal is somehow not sufficient to engage p97, and therefore proteasomal degradation fails.

**Figure 4.**
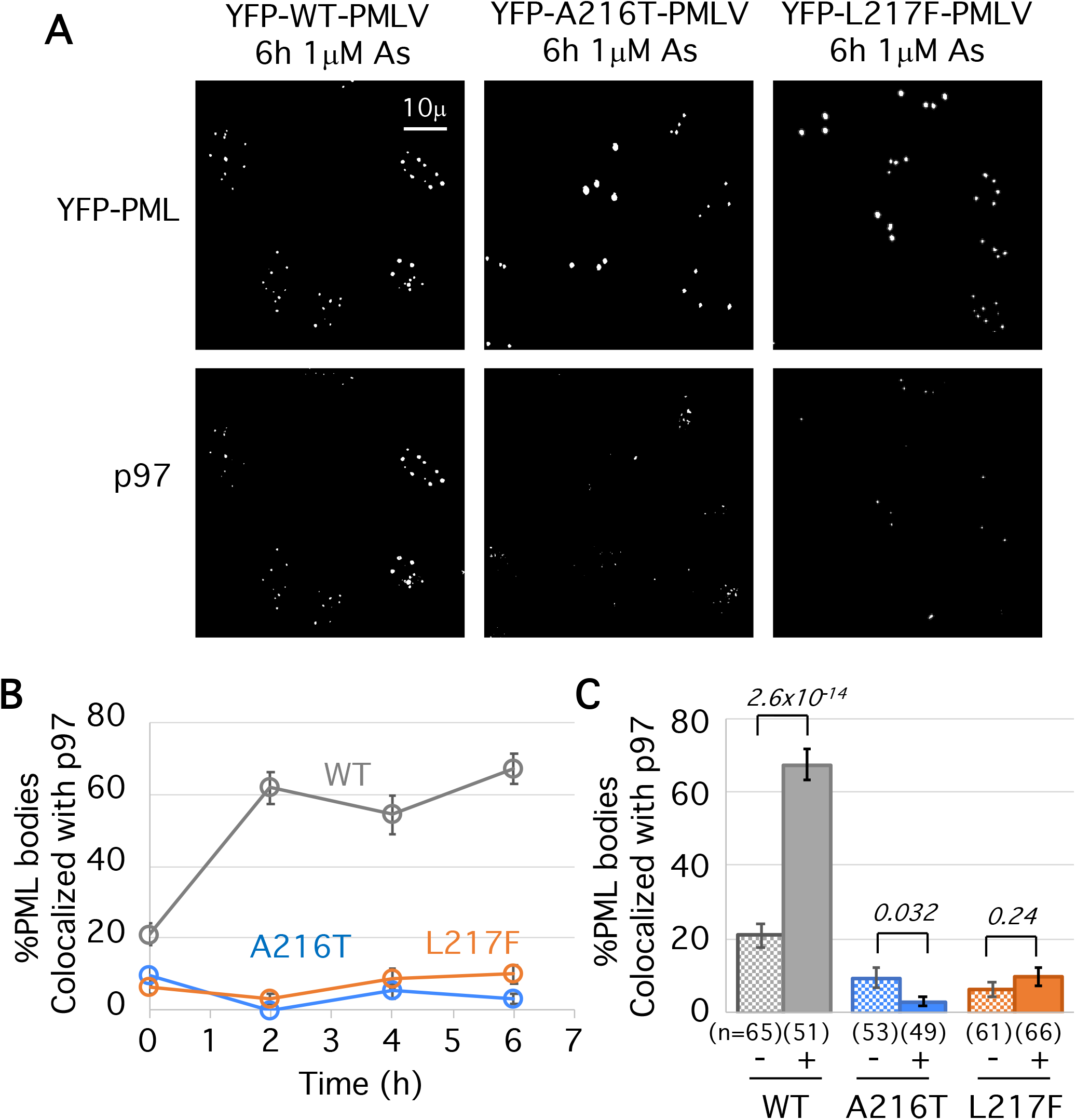
A216T and L217F mutants of PML do not recruit p97 in response to arsenic treatment. A. Immunofluorescence images showing YFP-PMLV and p97 in the indicated cell lines after 6 hours arsenic treatment. Data showing merged images for these and the 0h time-point data can be seen in supplementary figure 4. B. Quantitative summary of p97 colocalization with PML bodies in the indicated cell lines during 0h, 2h, 4h and 6h arsenic exposure. Markers show average values and error bars are standard error of the mean. C. Statistical analysis of the comparison between 0h and 6h arsenic exposure for PML-p97 colocalization. Columns represent average % colocalization, error bars are standard error of the mean, cell count is shown in brackets below each column and student’s two-tailed t-test p-values are shown above.

### Differential modification with SUMO and ubiquitin of PML mutants

To provide more detail on the post-translational modifications that accumulate on PML, we repeated the purification of YFP-PML nuclear bodies with a view to release SUMO or ubiquitin from resin-bound material by treatment with proteases specific for each (Supplementary Fig. 6A). The material released by protease treatment, and that remaining on the beads can be analysed by Western blotting, which should provide information not only on the covalent modification status of PML, but also that of the SUMO1, SUMO2/3 and ubiquitin species attached to PML (Supplementary Fig. 6B). Prior to purification, nuclear extracts for WT, A216T and L217F YFP-PMLV displayed the expected changes in electrophoretic mobility (Supplementary Fig. 6C), and were almost completely depleted from nuclear extracts after YFP-PML purification. Importantly, the modifications to PML were preserved during purification (Fig. 5A “No protease”). For all PML variants treatment of the beads with SENP1 returned most of the heavier PML forms back to more rapidly migrating species consistent with unmodified YFP-PMLV (Fig. 5 “SENP1” lanes). Treatment with USP2 had little impact on the distribution of PML species, indicating only a relatively small proportion of the modified PML is ubiquitin conjugated (Fig. 5A “USP2”). SUMO1 blots show that in untreated cells there are relatively low levels of SUMO1 associated with PML, while after arsenic treatment this increases dramatically for WT YFP-PMLV, modestly for L217F YFP-PMLV, and not at all for A216T YFP-PMLV (Fig. 5B). As expected, these species disappear upon SENP1 treatment, but are largely unaltered by treatment with USP2 (Fig. 5B). After arsenic exposure, the relative increase to SUMO2/3 modification is not as large as for SUMO1. Notably all three PML forms undergo an increase in SUMO2/3 conjugation, although A216T PML is modified to a much lesser degree than either WT or L217F (Fig. 5C). SUMO2/3 is also completely removed by SENP1, but USP2 treatment does not result in significant changes to SUMO2/3 antibody reactive species (Fig. 5C). Blotting with an antibody to ubiquitin reveals that it is present at low levels on WT, A216T and L217F YFP-PMLV in untreated cells but is dramatically increased on WT, modestly increased on L217F but any increase on A216T is undetectable in this assay (Fig. 5D). For both WT PML and the L217F mutant, SENP1 treatment not only reduces the amount of ubiquitin remaining on the beads, it also causes the ubiquitin-reactive species to run at a reduced molecular weight (Fig. 5D). This ubiquitin released by SENP1 is probably conjugated directly to SUMO, while the material remaining is directly conjugated to PML. As expected, ubiquitin is removed from purifications by treatment with USP2 (Fig. 5D). Although ubiquitination of L217F in response to arsenic is less than that of WT YFP-PMLV, it is still far more substantial than the A216T mutant (Fig. 5D).

**Figure 5.**
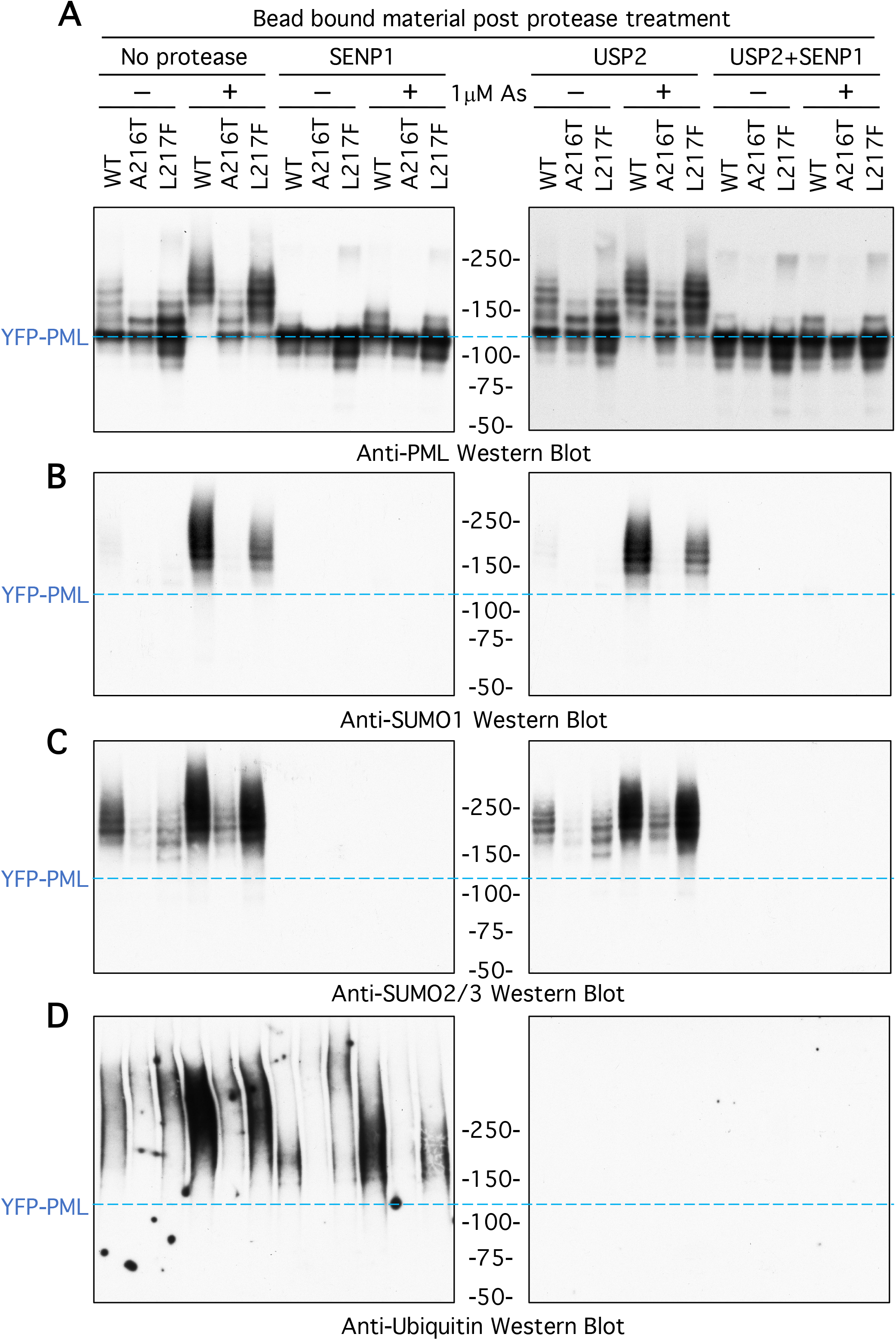
Failure of arsenic resistant mutants to generate the SUMO and ubiquitin modified intermediates required for degradation. PML bodies were enriched from U2OS PML-/- + YFP-PMLV WT, A216T or L217F cells, either untreated (-) or treated with 1μM arsenic (+) for 2 hr. Purified PML bodies bound to anti-GFP nanobody magnetic beads were either untreated (No protease), treated with SUMO specific protease (SENP1), treated with ubiquitin specific protease (USP2) or a combination of both (USP2 + SENP1). Material remaining bound to the beads after treatment was eluted and analysed by Western blotting using antibodies to A. PML, B. SUMO1, C. SUMO2/3 and D. ubiquitin.

The material released from the bead-associated YFP-PMLV by treatment with the proteases was also analysed by Western blotting. As expected, none of the treatments released YFP-PMLV (Supplementary Fig. 7A). SENP1 treatment released low levels of monomeric SUMO1 from untreated WT and L217F YFP-PMLV samples, with none detected from A216T purifications. After arsenic treatment the amount of SUMO1 released by SENP1 from WT samples increased for both WT and L217F PML forms, albeit less extensively for the mutant. Compared to these two, the amount of SUMO1 conjugated to A216T after arsenic exposure was negligible although it was at least detectable compared to the sample derived from untreated cells (Supp. Fig. 7B). SENP1 treatment of WT-PML purifications also released forms of SUMO1 shifted up around 10kDa from the unmodified SUMO1 species, which are presumed to be SUMO1 conjugated by ubiquitin, as they are sensitive to USP2 (Supplementary Fig. 7B). However, these species were not apparent in SENP1 elutions from L217F-PML purifications according to this assay, implying ubiquitination of SUMO1 is most extensive for in WT-PML. SENP1 digestion released substantial amounts of monomeric SUMO2/3 from preparations derived from untreated cells for all PML variants including A216T (Supp. Fig. 7C). Arsenic treatment induced a modest increase in SUMO2/3 conjugates on all three PML forms, although a ubiquitinated form of SUMO2/3 was only apparent for WT and L217F variants (Supp. Fig. 7C). These SUMO-ubiquitin hybrid species released by SENP1 were also detected using a ubiquitin antibody (Supplementary Fig. 7D). Although data shown in Fig. 5C suggest SUMO2/3 modification of PML is low prior to arsenic and increases after arsenic treatment, this is not consistent with analysis of the material released by SENP1 treatment (Supplementary Fig. 6C) which shows similar SUMO2/3 amounts both before and after arsenic. A possible explanation for this apparent discrepancy is that in untreated cells the SUMO2/3 modified PML are dispersed throughout the gel, whereas after SENP1 treatment all the SUMO2/3 runs as a single species.

In summary, A216T YFP-PMLV is defective for arsenic induced SUMO1, SUMO2/3 and ubiquitin modification. L217F YFP-PMLV appears to be modestly compromised for SUMO2/3 conjugation but shows much less than WT-PML levels of SUMO1 conjugation. Furthermore, WT PML shows higher levels of direct ubiquitination than L217F, whereas L217F ubiquitination appears to mostly take place on SUMO2/3 linked to PML. Finally, a significant proportion of SUMO1 conjugated to WT-PML is covalently conjugated by ubiquitin, while this was not detected for L217F PML.

### Quantitative proteomic analysis of PML post-translational modifications in response to arsenic

Early descriptions of PML modification by SUMO identified three major acceptors at lysines 65, 160 and 490 (Kamitani, Kito et al. 1998). More recently, the single largest SUMO site proteomic analysis to date (Hendriks, Lyon et al. 2017) described a total of 15 acceptor lysines in PML (Supplementary figure 8A). To provide quantitative data at the amino-acid site level for the post-translational modifications present in PML and SUMO, a peptide-level quantitative proteomics analysis was undertaken. Trypsin digestion leaves long SUMO adducts attached to modified lysines (Supplementary Fig 8B), that are challenging to identify by mass-spectrometry. Thus, a strategy combining GluC digestion and trypsin digestion was employed to give shorter peptide fragments. As GluC is less specific and less stable than trypsin, multiple GluC adducts from SUMOs remain on lysines after digestion (Supplementary Fig. 8B), which has the advantage of providing multiple peptide evidences for a single modification.

Cultures of U2OS PML-/- +YFP-PML cells were grown for the WT, A216T and L217F PML variants, and were either treated or not with 1μM arsenic for 2 hr (Supplementary Fig. 9A). Four experimental replicates for each condition were prepared and YFP-PML purified by the same method as described above. Purifications were analysed by SDS-PAGE (Supplementary Fig. 9B) and the uppermost section of the gel containing proteins larger than 100kDa, including YFP-PML, was processed for mass-spectrometry analysis. Anti-PML Western blot of a fraction of the inputs revealed the YFP-PML proteins responded as expected to arsenic treatment (Supplementary Fig. 9C). 21 phosphorylation, 14 SUMOylation and 11 ubiquitination sites were identified in PML (Supplementary datafile 1 and Supplementary Fig. 10A). Principal component analysis using intensity data for all modified and unmodified peptides from YFP-PML, SUMO1, SUMO2, SUMO3 and ubiquitin shows replicates cluster by experimental condition and are clearly separated from one-another (Supplementary Fig. 10B), indicating consistent differences among the conditions. Intensity data for all peptides were normalized by total YFP peptide intensity (Supplementary Fig. 10C), to minimize any systematic errors relating to PML expression or loading (Supplementary Fig. 10D). Using the total intensity of the unmodified peptides from SUMO1, SUMO2/3 and ubiquitin as a proxy for their overall protein abundance (Supplementary Fig. 10E-G), a remarkably similar picture to that revealed in the Western blot experiments above (Fig. 3) was found. Namely, prior to exposure to arsenic all three PML forms are associated with broadly similar amounts of SUMO1, SUMO2/3 and ubiquitin. Arsenic treatment has little effect on the modification of the A216T mutant, but triggers increased conjugation by SUMO1, SUMO2/3 and ubiquitin for WT and L217F PML, with the mutant showing a weaker response in SUMO1 and ubiquitin conjugation. Also consistent with the Western blot data is the finding that SUMO1 has a larger relative increase (5 fold) than SUMO2/3 (1.8 fold) in conjugation to WT PML after arsenic treatment. To more easily compare multiple protein/peptide intensities, each normalized intensity can be represented relative to the average intensity across all samples (Supplementary Fig. 10H), which can then be plotted in a single compact radial chart to show which differ the most between cell types and treatments (compare with Fig. 6A).

**Figure 6.**
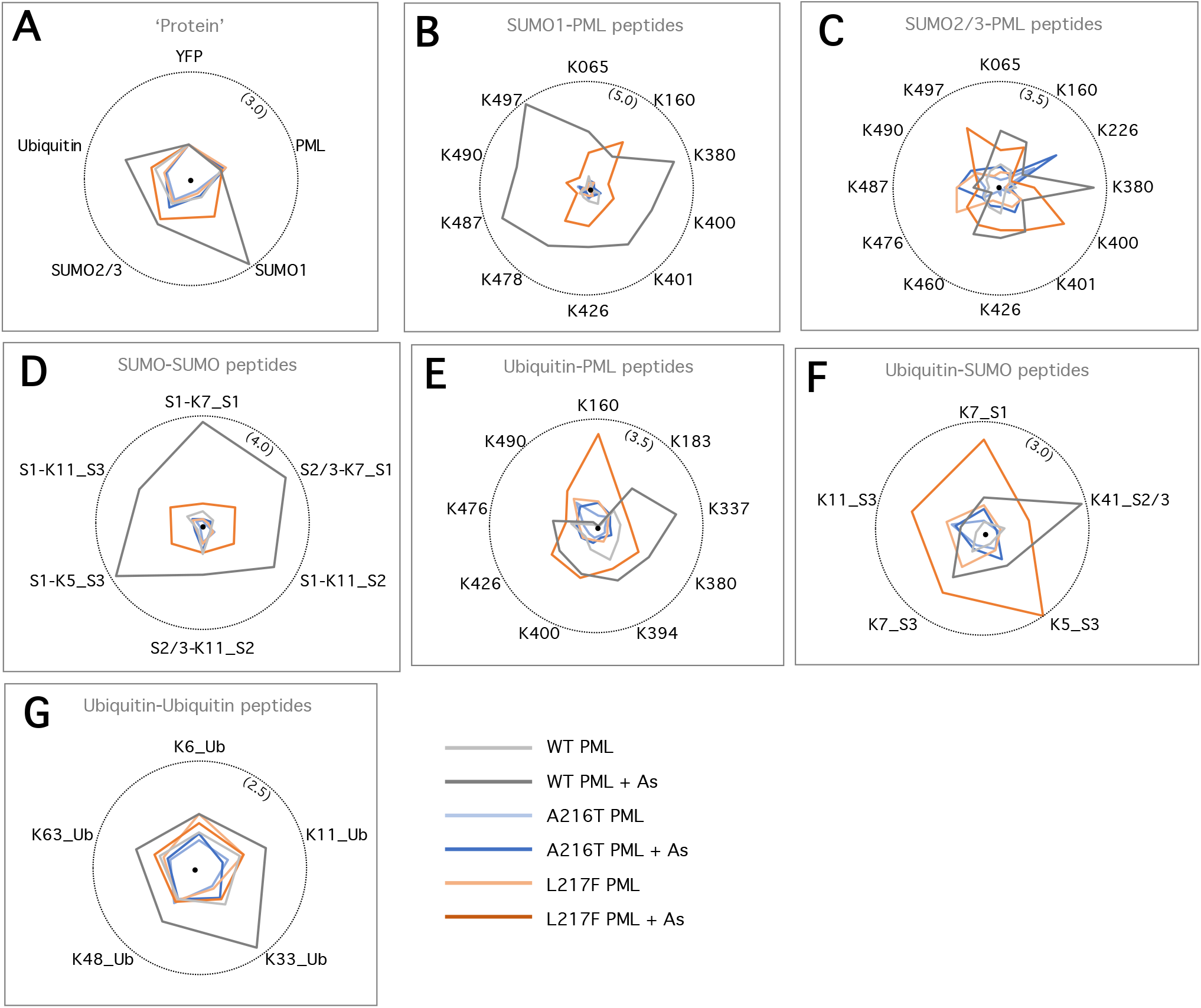
Differential site-specific modifications of WT, A216T and L217F PML forms before and after arsenic. PML bodies were enriched from U2OS PML-/- + YFP-PMLV WT, A216T or L217F cells and quantitatively analysed by mass spectrometry for protein content and site-specific modifications by SUMO1, SUMO2/3 and ubiquitin. A-G. Radial plots showing relative intensity for each of the indicated proteins or peptides. A. is derived from the total of the unmodified peptides for YFP, PML, SUMO1, SUMO2/3 and ubiquitin and is a proxy for of total protein abundance. Circles indicate the upper limit of the range, with the maximum relative intensity in brackets.

### Site-specific differences in SUMO modification to PML mutants

10 SUMO1 modifications to PML had reliable enough data for comparisons among conditions (Fig. 6B). After arsenic treatment, WT PML was much more extensively modified with SUMO1 than either mutant at all PML sites except K160 in B-Box1, which appeared to be more occupied by SUMO1 in the L217F mutant (Fig. 6B). Site- level analysis confirmed A216T PML was relatively unresponsive in SUMO1 conjugation. The results for SUMO2/3 were less clear-cut (Fig. 6C). Firstly, the scale of the difference among the three PML forms in the SUMO2 conjugation is much smaller than for SUMO1. Surprisingly, two of the 12 sites with good enough data for quantitative comparisons were most highly modified in the apparently inert A216T mutant (Fig. 6C). These were K226 close to the mutation hotspot and K487 in the NLS (Fig. 6C). Several other sites in or around the NLS, K400, K476, and K497, showed greater modification for L217F than WT PML (Fig. 6C). The WT protein only showed the most extensive SUMO2/3 modification at K380 (between the coiled-coil and NLS) and marginally higher conjugation at three sites; K65 (RING domain), K426 and K460 (both between the coiled-coil and the NLS). Site-level analysis of SUMO-SUMO branched peptides showed again that A216T PML was almost completely unresponsive (Fig. 6D). SUMO polymers did accumulate on L217F PML, but not to the same extent as for WT PML (Fig. 6D). Notably all bar one of the identified SUMO-SUMO linages contained SUMO1 as conjugate donor, acceptor, or both. Together these results show the higher levels of SUMO1 recruitment to WT PML compared to the mutants was via direct conjugation to PML, as well as SUMO1-SUMO polymer formation. Differences among the PML variants for SUMO2 conjuagation were more subtle and site-specific.

### Neither A216T nor L217F PML accumulate polyubiquitin chains in response to arsenic

Of the 11 ubiquitination sites identified in PML, the majority were sites also identified as SUMO acceptors, except K183 between B-Boxes 1 and 2 and K337 in the coiled-coil domain. 9 sites had high quality quantitative data which showed that A216T did not respond to arsenic with increased ubiquitination at any site (Fig. 6E). In general WT and L217F PML showed similar global increases in ubiquitination on PML although there were differences at the site level. Notably both A216T and L217F PML were much more highly modified at lysines 160 and 490 than WT (Fig. 6E). These are two of the three sites originally identified as major SUMO acceptors, and likely to already have relatively high occupancy with SUMO. K183 and K337 were more occupied by ubiquitin in WT PML than either mutant (Fig. 6E). There is a striking difference between WT and L217F in the ubiquitination of the associated SUMO (Fig. 6F). 4 of the 5 ubiquitination sites in SUMO 1 and SUMO2/3 quantified were more extensively ubiquitinated in L217F PML purifications then the WT equivalent (Fig. 6F). Conversely, ubiquitin-ubiquitin conjugations were generally more abundant in WT PML purifications than for L217F PML (Fig. 6G). These results show that compared with WT PML, arsenic-induced ubiquitination of L217F PML is skewed towards SUMO ubiquitination, and away from ubiquitin-ubiquitin polymer formation.

### SUMO1 is essential for arsenic-induced PML degradation

The finding that the stable L217F PML mutant is more defective in SUMO1 conjugation than SUMO2 is unexpected. RNF4 has been shown to have specificity for polymeric SUMO conjugates, with a chain of 4 or more SUMOs having higher affinity for RNF4 than shorter polymers or monomeric SUMO conjugates (Tatham, Geoffroy et al. 2008). Due to the presence of an internal SUMO conjugation consensus motif in SUMO2/3 but not SUMO1, it is assumed that SUMO2/3 form branched conjugates more readily than SUMO1, and therefore represent the more common RNF4 substrate. However, SUMO1 can still participate in polySUMO conjugates as a SUMO donor, and proteomics studies have also found SUMO1 to be an acceptor for SUMOylation, although the salience of this is poorly understood. To investigate the role of SUMO1 in arsenic-induced PML degradation, a U2OS cell line lacking SUMO1 (SUMO1 -/- U2OS) was generated by CRISPR-cas9 genome editing. Immunofluorescence confirmed the loss of SUMO1 expression in these cells (Fig. 7A). Anti-PML Western-blot analysis of crude cell extracts of cells exposed to 1µM As over 24 hr showed that in cells lacking SUMO1 there is a molecular weight shift in PML in a manner similar to SUMO1 +/+ cells (Fig. 7B), but the modified forms of PML appear to be more resistant to degradation. These persist up to 24 hr after addition of As to the medium, when PML from WT cells is almost completely degraded (Fig. 7B). Furthermore, after these extended periods of As exposure, unconjugated PML begins to re-accumulate in SUMO1 - / cells (Fig. 7C), almost returning the cellular pool of PML to the initial state (Compare 0h and 24h lanes in Fig. 7C). This confirms SUMO1 is essential for PML degradation and insufficient conjugation of the L217F PML mutant by SUMO1 potentially explains its stability during arsenic exposure.

**Figure 7.**
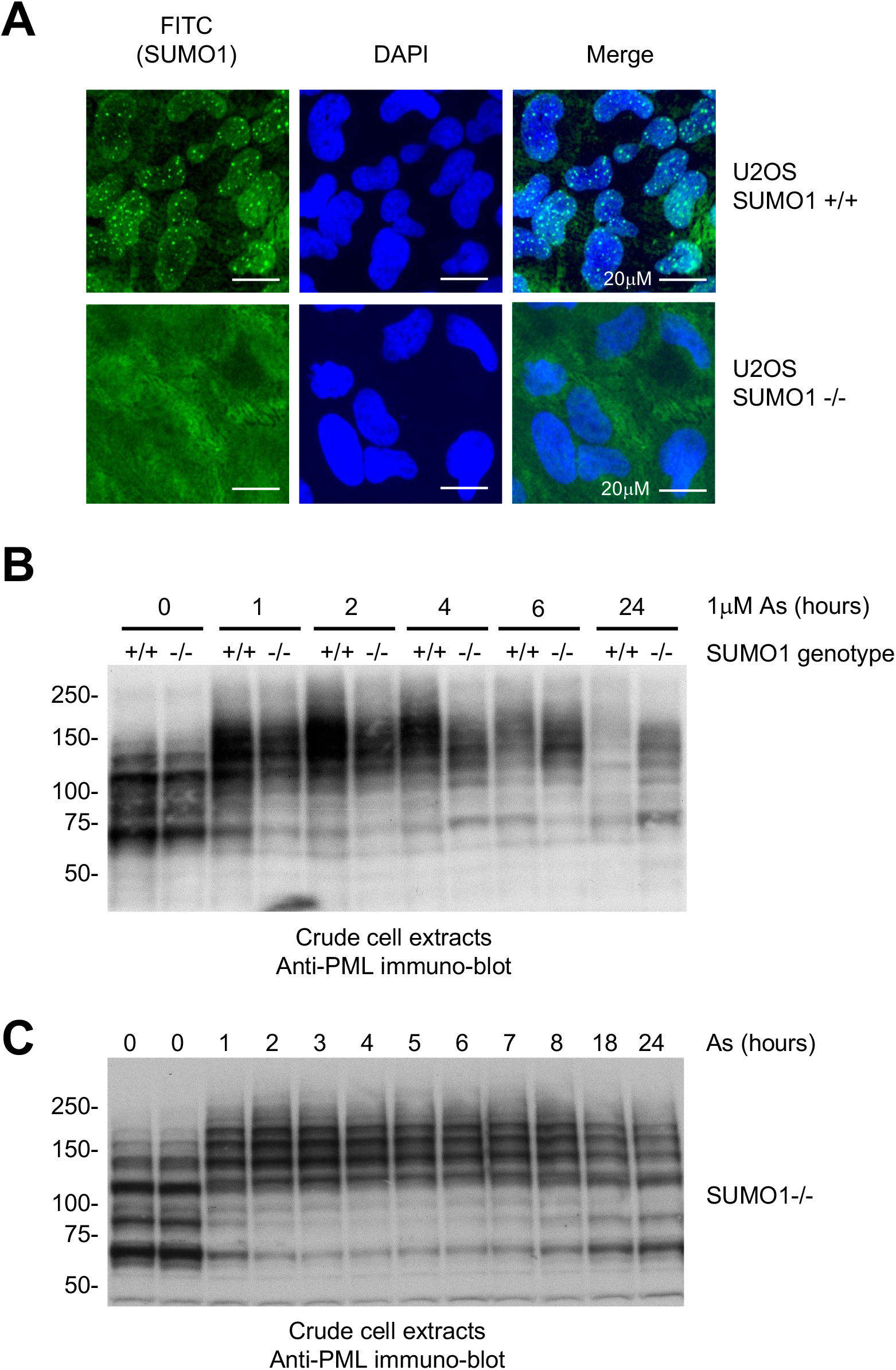
Effect of SUMO1 deletion on arsenic-induced WT PML degradation. A. Wild-type (SUMO1 +/+) and SUMO1 knockout (SUMO1 -/-) U2OS cells were exposed to 1µM As for 2 hours to allow SUMO1 recruitment to PML bodies, fixed and double stained with DAPI and anti-SUMO1 (FITC) before image acquisition using InCell analyser 2200 high-content imaging microscope and differential image visualization by ImageJ software. B. SUMO1 +/+ and SUMO1 -/- U2OS cells were exposed to 1µM As for the indicated periods before lysis and analysis by Western blot for PML. C. As in B except only for SUMO -/- cells and with more time-points.

## DISCUSSION

The overarching cellular mechanisms governing arsenic-induced degradation of PML and the oncogenic fusion PML-RARA were broadly characterised more than ten years ago (Lallemand-Breitenbach, Jeanne et al. 2008, Tatham, Geoffroy et al. 2008). Arsenic triggers the SUMOylation of PML, followed by its ubiquitination, then degradation by the proteasome. However, the precise molecular signals and effector proteins required to progress through each step are not fully understood. The functional characterisation of mutant forms of PML and PML-RAR found in APL patients refractory to arsenic treatment, provides an opportunity to better understand the biochemical details of arsenic-induced PML degradation by understanding their loss of function. We have used a combination of high content, time-lapse microscopy, and peptide-level proteomic analysis to study two common patient- derived PML mutants; A216T and L217F. We generated three model U2OS cell lines lacking endogenous PML but expressing YFP fusions of WT PML, A216T PML or L217F PML. In untreated cells the two mutants of PML formed atypical nuclear bodies, fewer in number and larger in size than WT bodies, and both mutants were stable when cells were exposed to 1μM arsenic for 24 hours. Importantly, during arsenic treatment the modification status of each was quite different, and both differed from WT PML.

After 2 hours exposure to 1μM arsenic, A216T PML appeared almost inert and showed little increase in either SUMOylation or ubiquitination. Thus, A216T PML stability is explained by a failed SUMOylation response early in the degradation pathway. In contrast, arsenic did trigger SUMO and ubiquitin conjugation to L217F PML, and yet it was not degraded. Site level proteomics revealed that while ubiquitination did increase for L217F PML, it differed from WT PML in the precise lysine sites to which it conjugated. The ubiquitin associated with L217F appeared to favour attachment to SUMO molecules, while on WT-PML ubiquitin accumulated more in polyubiquitin chains. This lack of polyubiquitin chain formation potentially explains the observed lack of recruitment to L217F-PML of the segregase p97, which recognises ubiquitin chains and is essential for PML degradation (Jaffray, Tatham et al. 2023). p97 recognises polyubiquitin chains containing 5 or more copies of ubiquitin (Twomey, Ji et al. 2019), and both K11 and K48 linkages are implicated in p97 function (Locke, Toth et al. 2014, Meyer and Rape 2014, Yau, Doerner et al. 2017). Crucially, WT-PML showed statistically significant increases in ubiquitin- ubiquitin linkages via K11, K33 and K48 (Supp. Fig. 11A). Although these increases were modest, we expect an appropriate degradation signal for PML to be short-lived in cells, making their detection difficult in studies like these.

Arsenic exposure triggered a much greater SUMO1 response on WT PML than it did for SUMO2/3. From these studies we cannot determine whether more SUMO1 than SUMO2/3 was conjugated to PML, but relative to initial conjugation levels, the SUMO1 response was much greater than SUMO2/3. Furthermore, the L217F mutant displayed lower levels of SUMO1 conjugation upon arsenic exposure than WT PML, while total amounts of SUMO2/3 associated with both PML forms were comparable. The importance of SUMO1 was further evidenced by the finding that WT PML is stable in cells lacking SUMO1, suggesting that SUMO2/3 cannot compensate for its loss. This raises the possibility that sub-optimal SUMO1 conjugation may lead to the failure of L217F to generate the appropriate ubiquitin signal for p97 binding.

Based on the current model for PML degradation, critical requirement for SUMO1 seems counter intuitive. The E3 ligase responsible for PML ubiquitination, RNF4 is poly-SUMO specific (Tatham, Geoffroy et al. 2008), and by lacking an internal SUMO conjugation consensus, SUMO1 is thought not to form polySUMO chains as readily as SUMO2/3. Incorporation of SUMO1 into SUMO2/3 polymers could suppress further polymer formation by blocking lysines in SUMO2/3, and so SUMO1 may be expected to be inhibitory to PML degradation via RNF4. One potential explanation is enough SUMOs in close proximity could substitute for a SUMO polymer, and engage and activate RNF4 (Aguilar-Martinez and Sharrocks 2016). This may be more likely for a protein like PML which forms multimeric complexes and contains numerous SUMOylation sites. It is also conceivable that independent of RNF4, SUMO1 provides a unique signal recognised by an as-yet unidentified protein effector of the PML degradation pathway that specifically recognises SUMO1 and not SUMO2/3. These prospects will require further investigation. Alternatively, rather than generating a new signal, SUMO-1 may function to suppress the formation of unproductive signals. For example SUMO1 may compete with ubiquitin for conjugation to lysine residues on proteins other than ubiquitin itself, and so promote polyubiquitin chain formation. In this way SUMO1 modification of lysines from SUMOs attached to WT-PML or PML itself may block spurious ubiquitination and effectively drive the synthesis of ubiquitin-ubiquitin linkages required for degradation. L217F PML which is less SUMO1 modified than WT-PML might then have more available non-ubiquitin lysines and so does not synthesise ubiquitin chains as readily. Consistent with this, ubiquitination of SUMOs was found to be generally higher for L217F PML than WT (Fig. 6F), and SUMO1-SUMO linkages were more abundant in WT-PML preps than L217F (Fig. 6D).

In summary, two patient-derived mutant forms of PML, A216T and L217F result in similar clinical aetiologies but have subtly different biochemical responses to arsenic resulting in blocked degradation (See Supplementary figure 12). A216T is almost completely unresponsive to arsenic, experiencing only very modest changes in SUMOylation and ubiquitination, most likely because it does not engage the appropriate SUMO conjugation machinery. L217F PML does become modified with both SUMO1 and SUMO2/3, but while ubiquitination is stimulated, it does not create the appropriate signal required to recruit p97 prior to degradation by the proteasome. These results reveal the subtle intricacies of the PML degradation pathway, and propose a higher level of complexity than previously considered.

## MATERIALS AND METHODS

### Preparation of anti-GFP nanobody magnetic beads

300mg of Dynabeads M-270 Epoxy (ThermoFisher Scientific Catalogue No. 14302D) were resuspended in 10 ml of 0.1 M sodium phosphate buffer pH 7.5 and washed twice using a DynaMag-15 magnet (ThermoFisher Scientific Catalogue No. 12301D) with 10 ml of the same buffer. 4 mg of the LaG16 nanobody that recognises GFP (Fridy, Li et al. 2014) in 10 ml of 1 M (NH_4_)_2_SO_4_, 0.1 M sodium phosphate buffer pH 8.0 was added to the beads which were incubated overnight at 27°C with end over end rotation. After coupling was complete the beads were washed 6 times with phosphate buffered saline (PBS) and stored in PBS, 0.1% NaN_3_ at 4^0^C.

### Purification of YFP-PML from U2OS PML-/- YFP-PML cells

For each ‘replicate’ of the proteomics experiment five 15 cm diameter plates of approximately 80% confluent cells were used. Prior to harvesting, growth medium was removed and cells washed in situ twice with 5mL PBS/100mM iodoacetamide. For each dish, cells were scraped into 5mL PBS+100mM iodoacetamide and transferred to a 50mL tube. A second 2mL volume of PBS+100mM iodoacetamide was used to clean the plate of all cells and pooled with the first 5mL. This was repeated for all dishes for each replicate and pooled in the same 50mL tube (∼35mL total). Cells were pelleted by centrifugation at 400g for 5 minutes at 22°C and resuspended in 10 mL 4°C hypotonic buffer including iodoacetamide (10mM HEPES pH 7.9, 1.5mM MgCl2, 10mM KCl, 0.08% NP-40, 1x EDTA-free protease inhibitor cocktail (Roche), + 100mM Iodoacetamide). A sample of this may be retained as a “Whole cell extract”. These were snap-frozen in liquid nitrogen and stored at −80°C until required. Cells were thawed by tube rotation at 4°C for 30 minutes. Tubes were then centrifuged at 2000g, 4°C for 10 minutes to pellet nuclei. Supernatants were discarded but a sample of this can be taken as a “Cytosolic fraction”. Each pellet of nuclei was resuspended in 5mL ice-cold hypotonic buffer containing 20mM DTT but without iodoacetamide, transferred to a 15mL tube and centrifuged at 2000g, 4°C for 10 minutes. The nuclei were washed once more in 5mL ice-cold hypotonic buffer (without iodoacetamide or DTT) and the supernatant discarded. To digest DNA, nuclei were resuspended in 5mL “Buffer A” (50mM Tris pH 7.5, 150mM NaCl, 0.2mM DTT, 1µg/ml benzonase, 1x EDTA-free protease inhibitor cocktail (Roche)) and incubated on ice for 15 minutes. Samples were sonicated in 15mL tubes on ice (Branson Sonifier) using the small probe, 6 cycles of 30 seconds at approximately 50% amplitude with at least 2 minutes cooling on ice in between cycles. PML bodies should remain intact at this point. Insoluble debris was removed by centrifugation at 1200g for 20 minutes at 4°C, and the supernatant containing soluble PML bodies, transferred to a new 15ml centrifuge tube. A sample of this can be taken as a “Nuclear fraction”. To each 5mL supernatant 6mL “Buffer B” was added (50mM Tris pH 7.5, 1% Triton-X-100, 0.1% Deoxycholic acid, 1x EDTA-free protease inhibitor cocktail (Roche)). To each 11mL solution, 1mL volume of a 5% beads:buffer slurry (v:v) of magnetic anti-GFP nanobody beads, pre-equilibrated in 1:1.4 Buffer A:B mix (v:v), was added and mixed on a tube roller at 4°C for 16h (50μL beads per sample). Using the DynaMag rack (Thermofisher scientific) the beads were extracted from suspension and the supernatant removed. A sample of this can be analysed to ensure YFP-PML depletion. Beads were then washed with 5mL “Buffer C” (50mM Tris pH 7.5, 500mM NaCl, 0.2mM DTT) and the beads extracted. These beads were then resuspended in 1mL “Buffer D” (50mM Tris pH 7.5, 150mM NaCl, 0.2mM DTT) and transferred to a new 1.5mL protein LoBind tube (Eppendorf) and after bead separation, washed once more with 1mL Buffer D. One half of this final sample of beads (25μL beads) was eluted for proteomic analysis by adding 35µl 1.2X NuPAGE LDS sample buffer (Thermofisher scientific), and incubated at 70°C for 15 minutes with agitation, followed by fractionation by SDS-PAGE. If Western blot analysis was also required, the second half of the resin (25μL) would be resuspended in 250µl 1.2X NuPAGE LDS sample buffer (Thermo) and incubated at 70°C for 15 minutes with agitation. Approximately 30µl would be used per gel lane. If treatment with proteases was required to monitor PML modification status, then the second half volume of beads (25μL) was divided into four and treated as follows: 1-Add 150 µl Buffer D. 2-Add 150 µl 250nM SENP1 in Buffer D. 3- Add 150 µl 500nM USP2 in Buffer D. 4- Add 150 µl 250nM SENP1/500nM USP2 in Buffer D. All tubes were agitated for 1-2 hrs at 22°C, supernatants removed (and can be kept for analysis) and beads washed twice with 1mL buffer D before elution with 250 µl 1.2X LDS, 70°C, 15 minutes with agitation.

### Proteomic analysis of YFP-PML purifications

Elutions from anti-GFP nanobody beads in 1.2x LDS/60mM DTT protein sample buffer were fractionated on 4-12% polyacrylamide Bis-Tris NuPAGE gels (Thermofisher Scientific) using MOPS running buffer. After Coomassie staining and destaining gels were excised into 4 slices per lane and tryptic peptides extracted from top slice (>100kDa) as previously described (Shevchenko, Tomas et al. 2006). Peptides were resuspended in 40μL 0.1% TFA 0.5% acetic acid and half STAGE tip purified prior to digestion with GluC. This gave two fractions per replicate: Trypsin digested, and Trypsin+GluC digested. 6μL of each was analysed by LC-MS/MS. This was performed using a Q Exactive mass spectrometer (Thermofisher Scientific) coupled to an EASY-nLC 1000 liquid chromatography system (Thermofisher Scientific), using an EASY-Spray ion source (Thermofisher Scientific) running a 75 μm x 500 mm EASY-Spray column at 45°C. A 150 minute elution gradient with a top 8 data-dependent method was applied. Full scan spectra (m/z 300–1800) were acquired with resolution R = 70,000 at m/z 200 (after accumulation to a target value of 1,000,000 ions with maximum injection time of 20 ms). The 8 most intense ions were fragmented by HCD and measured with a resolution of R = 17,500 at m/z 200 (target value of 500,000 ions and maximum injection time of 120 ms) and intensity threshold of 2.1×10^4^. Peptide match was set to ‘preferred’, a 40 second dynamic exclusion list was applied and ions were ignored if they had unassigned charge state 1, 8 or >8.

MS data were processed in MaxQuant version 1.6.1.0 (Cox and Mann 2008, Cox, Neuhauser et al. 2011). Multiple MaxQuant runs were performed considering multiple SUMO1 and SUMO2/3 derived adducts as variable modifications (see Supplementary Datafile 1 for details), as well as those typical for ubiquitin and phosphorylation to Ser and Thr residues. A MaxQuant search for SUMO adducts was also conducted using a concatenated database approach using linear peptide fusions of SUMO C-termini to substrate peptides as search library (Matic, van Hagen et al. 2008). Oxidised methionine and acetylated protein N-termini were also selected as variable modifications and carbamidomethyl-C was the only fixed modification. Maximum number of variable modifications was set to 4. The match between runs option was enabled, which matches identified peaks among slices from the same position in the gel as well as one slice higher or lower. The uniport human proteome database (downloaded 19/4/2019 - 73920 entries) digested with Trypsin/P (maximum missed cleavages was 5) was used as search space. LFQ intensities were required for each slice. All FDR filtering was set to 1%. For peptides derived from GluC digestions 8 maximum missed cleavages were considered. Raw peptide intensities were manually normalized by the total YFP peptide intensity for each sample. Mass spectrometry data from samples derived from the same replicate were numerically summed to provide a final normalized peptide intensity for each identification in each sample. Downstream data processing used Perseus v1.6.1.1 (Tyanova, Temu et al. 2016). Zero intensity values were replaced from log_2_ transformed data (width 0.3 and downshift 1.8).

### Pre-extraction of cells

Cells were grown on coverslips in 24 well plates. Coverslips were washed twice with PBS before being exposed to pre-extraction buffer (25 mM HEPES pH 7.4, 50 mM NaCl, 3 mM MgCl2, 0.5% Triton-X-100, 0.3M Sucrose) for 10 min. Cells were then processed through the procedure below for immunofluorescence.

### Immunofluorescence analysis

Adherent cells were washed three times with PBS, fixed with 4% Formaldehyde in PBS for 15mins, then washed again three times with PBS before blocking for 30 min in 5% BSA, 0.1% TWEEN 20 in PBS. Fixed cells were washed once with 1% BSA, 0.1% TWEEN 20 in PBS before incubation for 1-2 hr with primary antibody in 1% BSA, 0.1% TWEEN 20 in PBS. These were chicken anti-PML (in house) and sheep anti-p97 (MRC Dundee). After 3 washes in 1% BSA, 0.1% TWEEN 20 in PBS cells were incubated with secondary antibody in 1% BSA, 0.1% TWEEN 20 in PBS for 60 min., washed three times in 1% BSA, 0.1% TWEEN 20 in PBS and stained with 0.1µg/ml DAPI for 2 min. These were Alexa Fluor (Invitrogen) Donkey anti-chicken 594, donkey anti-sheep 594 and donkey anti-chicken 488. Cover slips were rinsed twice with PBS, twice with water and dried before mounting in MOWIOL. Images were collected using a DeltaVision DV3 widefield microscope and processed using Softworx (both Applied Precision, Issaquah, WA). Images are presented as maximal intensity projections using Omero software. For quantitative p97-PML colocalization analysis the same image settings were applied across all fields and PML bodes per cell were counted for between 20 and 40 cells per condition. P97 colocalisation was defined as any p97 signal above background within the same region as a PML body. Percent colocalization was then calculated for each cell.

### Live-Cell Imaging in Real Time and Measurement of Fluorescence Intensities

For all live-cell imaging experiments, cells were seeded onto μ-slide 8 well glass bottom (Ibidi 80827). Immediately prior to imaging cell medium was replaced with DMEM minus phenol red (gibco 31053-038) supplemented with 2mM glutamine and 10% fetal bovine serum. Cells were untreated or treated with 1μM arsenic. Time-laspe microscopy was performed on a DeltaVision Elite restoration microscope (Cytiva) fitted with an incubation chamber (Solent Scientific) set to 37 °C and 5% CO2 and a cooled charge-coupled device camera (CoolSnap HQ; Roper). SoftWorx software was used for image collection. Datasets were deconvolved using the constrained iterative algorithm (Swedlow, Sedat et al. 1996, Wallace, Schaefer et al. 2001) using SoftWorx software. Using a YFP specific filter set (Ex510/10nm, Em537/26nm), 5 z-planes spaced at 2 µm were taken every 15 min. A single brightfield reference image was taken at each time point to monitor cell health. Time courses were presented as maximum intensity projections of deconvolved 3D datasets. Movies were created in OMERO using the deconvolved images.

### Quantitative analysis of PML body composition during Live-Cell imaging

Batch analysis of time-lapse movies was carried out using ImageJ (Schneider, Rasband et al. 2012) macros in Fiji (Schindelin, Arganda-Carreras et al. 2012) (Supplementary Materials). Briefly: 1) datasets (deconvolved with SoftWoRx) were cropped to remove border artefacts, 2) for each timepoint the maximum focus slice was selected according to intensity variance (radius 2 pixels), 3) auto-thresholding was performed using a threshold of mean intensity + 3 standard deviations, and 4) ImageJ’s built-in “Analyze Particles” function was used to measure PML body count, size, total area, and intensity. Results were aggregated and summarized using R scripts as described previously (Jaffray, Tatham et al. 2023).

### Preparation of cell extracts and Western blotting

Adherent cells were washed with PBS and lysed by the addition of 1.2X LDS buffer and heated to 70°C for 10 min. Cell lysates were sonicated using a probe sonicator for two bursts of 30 sec. Samples were run on NuPage 3-8% or 4-12% Bis-Tris precast gels (ThermoFisher Scientific) and transferred to nitrocellulose. Membranes were blocked in PBS with 5% non-fat milk and 0.1% Tween-20. Primary antibody incubations were performed in PBS with 3% BSA and 0.1% Tween-20. For the HRP-coupled secondary antibody incubations 5% milk was used instead of the BSA. The signal was detected by Pierce enhanced chemiluminescence (ThermoFisher scientific 32106) and X-ray films. Antibodies. Chicken anti-PML (homemade), sheep anti-SUMO-1 (homemade), rabbit anti-SUMO- 2 (Cell signaling 4971S), rabbit FK2 ubiquitin (Enzo BML-PW8810-0500), rabbit anti-p97 (Abcam ab109240). HRP-coupled secondary anti-mouse, anti-rabbit anti-chicken and anti-sheep were purchased from Sigma.

## Supporting information

Supp. Datafile 1

---

**Supplementary Figure 1.**
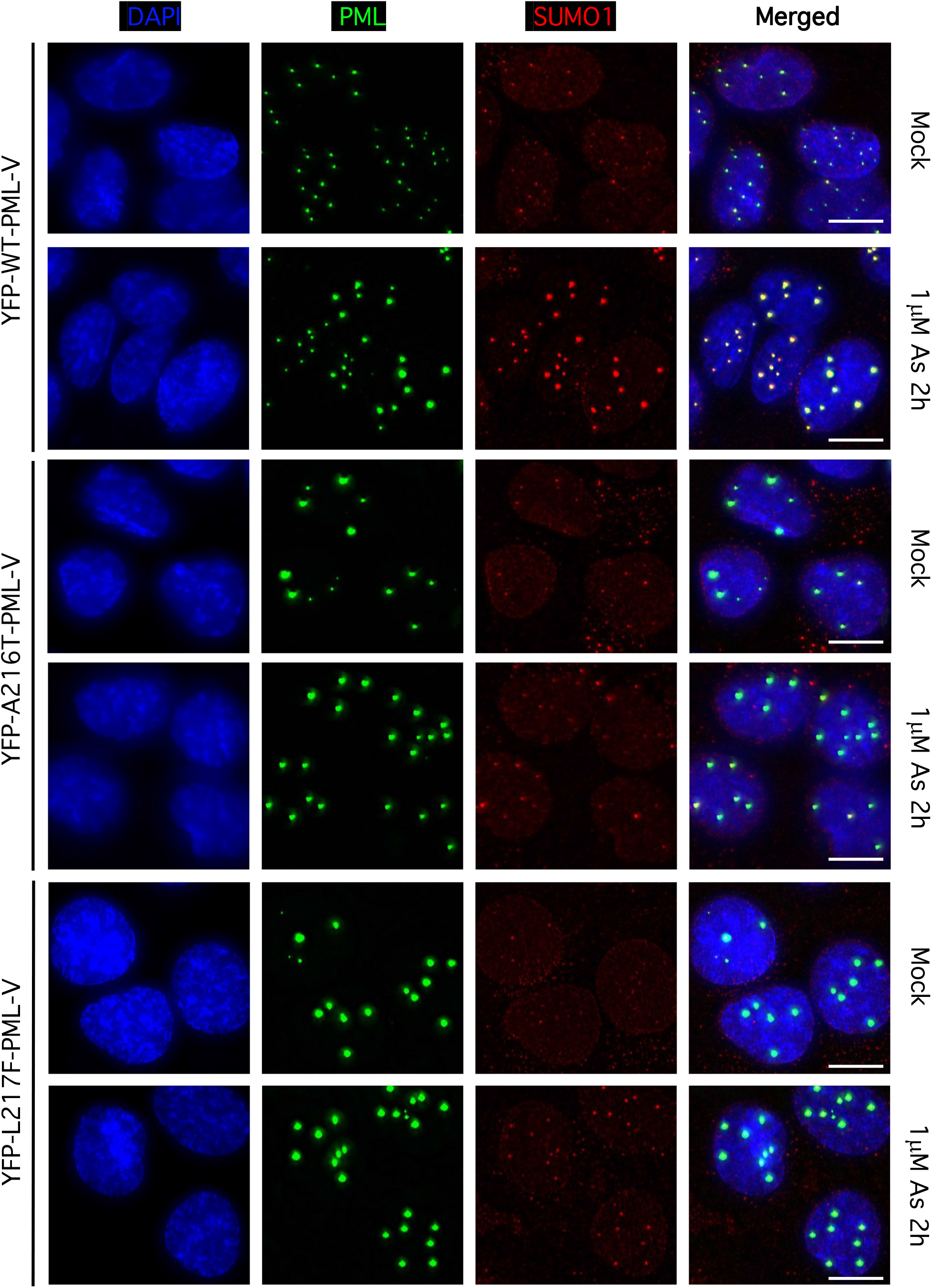
Arsenic-induced colocalisation of SUMO1 with WT, A216T and L217F forms of YFP-PML. Colocalisation of SUMO1 with WT, A216T and L217F PML variants after 1μM arsenic treatment for 2h studied by fluorescence microscopy. Scale bar is 10μm.

**Supplementary Figure 2.**
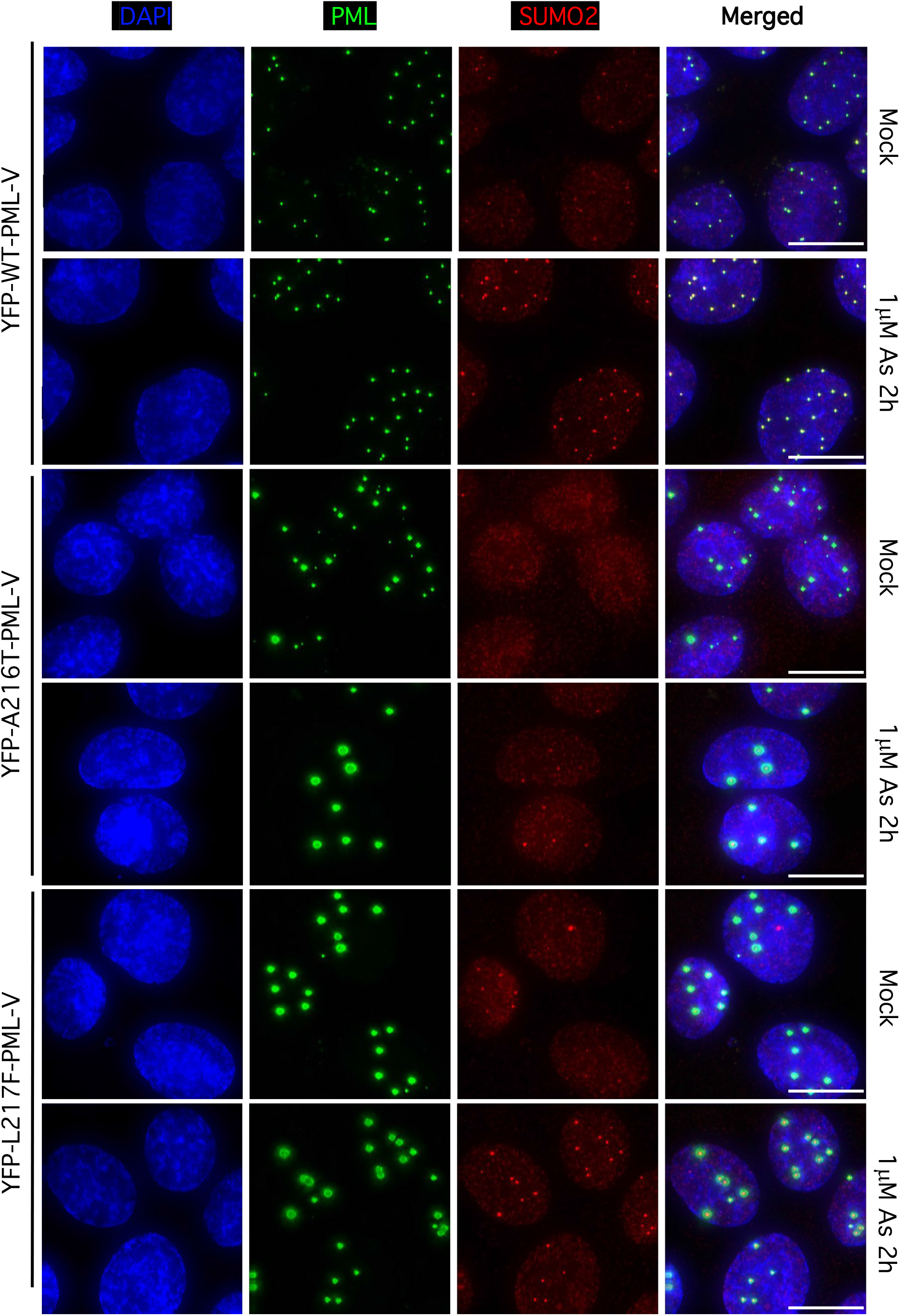
Arsenic-induced SUMO2/3 colocalisation with WT, A216T and L217F forms of YFP-PML. Colocalisation of SUMO2/3 with WT, A216T and L217F PML variants after 1μM arsenic treatment for 2h studied by fluorescence microscopy. Scale bar is 10μm.

**Supplementary Figure 3.**
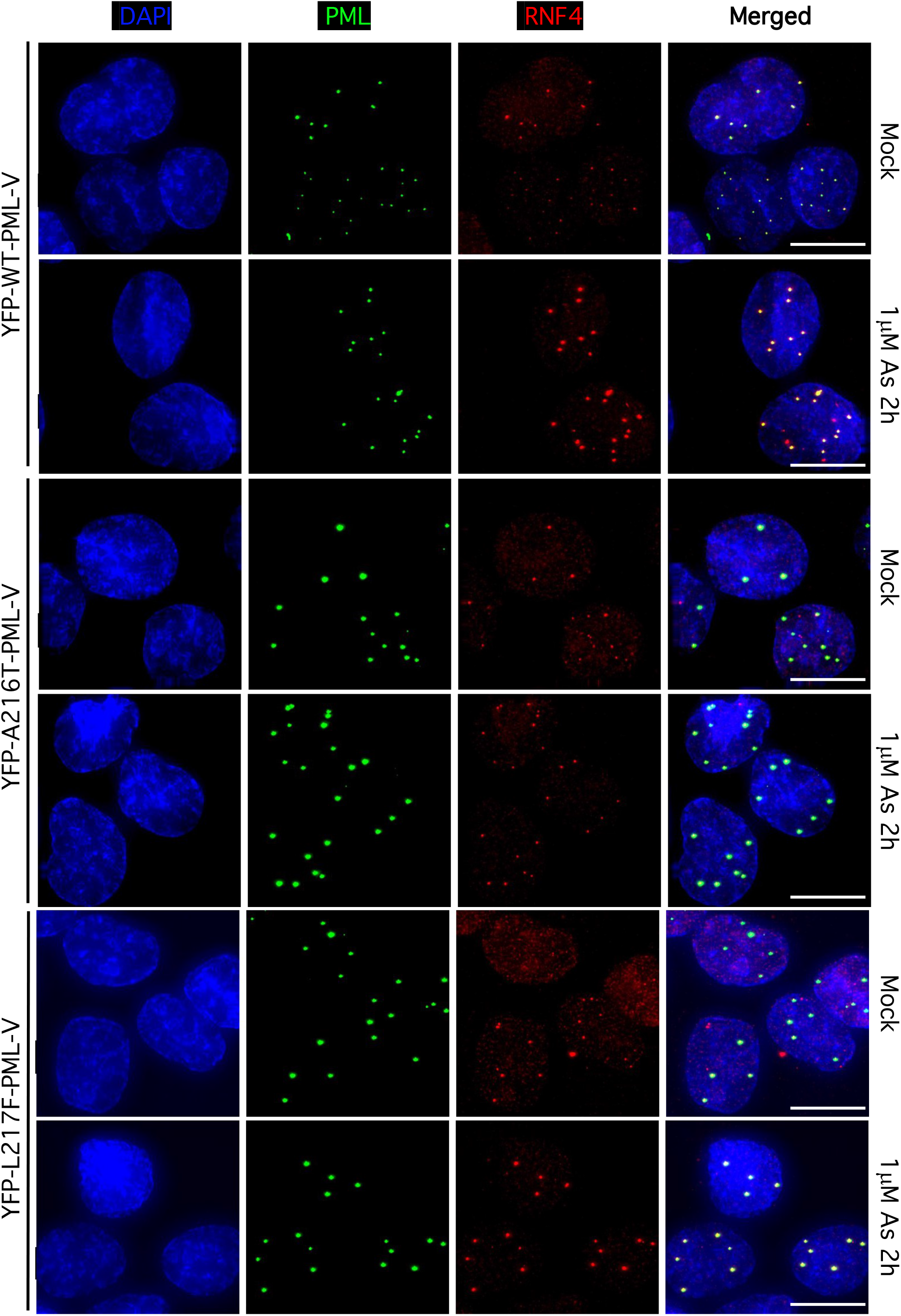
Arsenic-induced RNF4 colocalisation with WT, A216T and L217F forms of YFP-PML. Colocalisation of RNF4 with WT, A216T and L217F PML variants after 1μM arsenic treatment for 2h studied by fluorescence microscopy. Scale bar is 10μm.

**Supplementary Figure 4.**
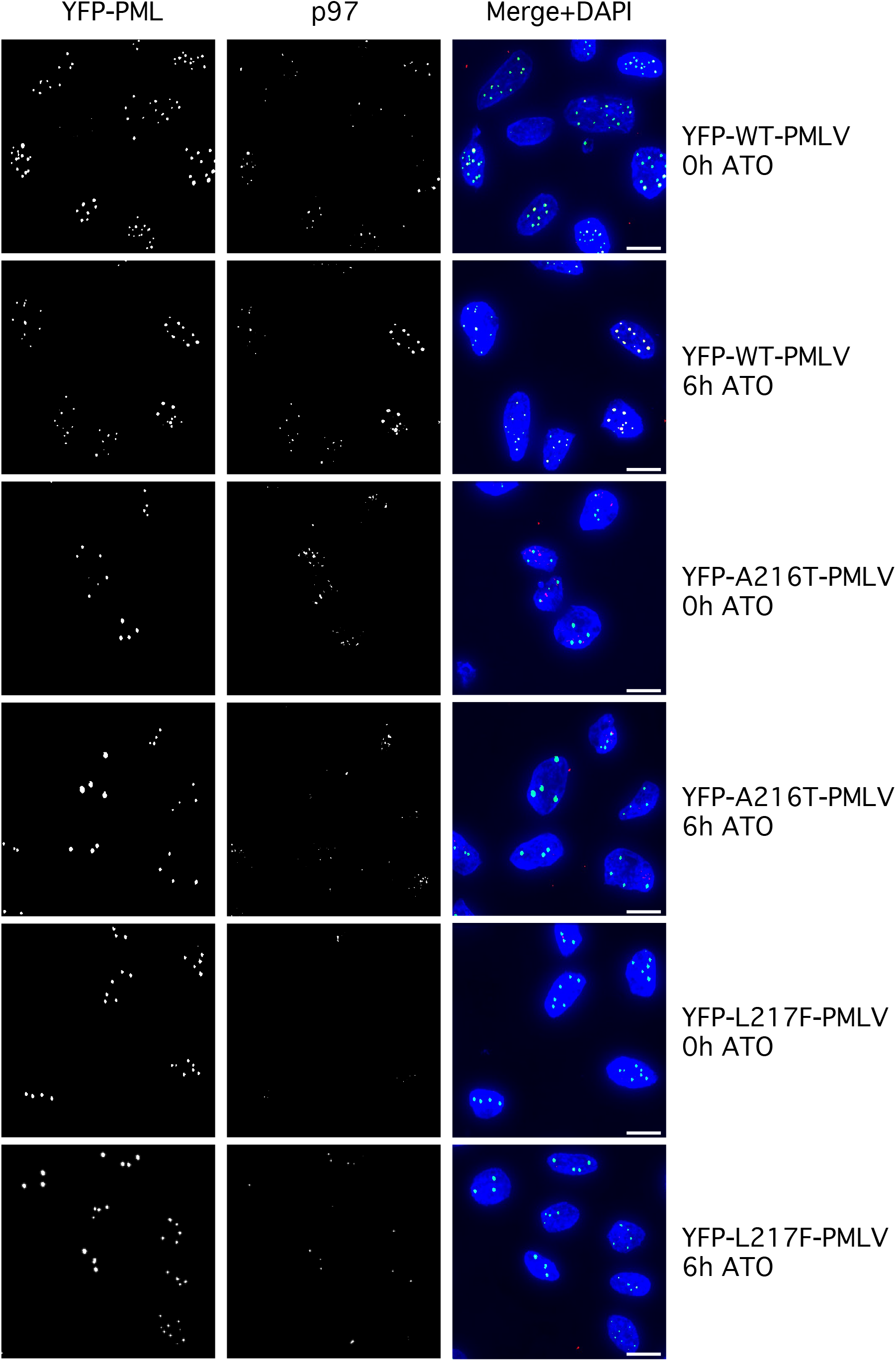
Arsenic-induced p97 colocalisation with WT, A216T and L217F forms of YFP-PML. Representative immunofluorescence images for 1μM arsenic treated YFP-PMLV cells at 0 hr and 6 hr. Monochrome images for YFP-PML and p97 channels are shown along with the merge of YFP-PMLV (green) and p97 (red) with DAPI (blue).

**Supplementary Figure 5.**
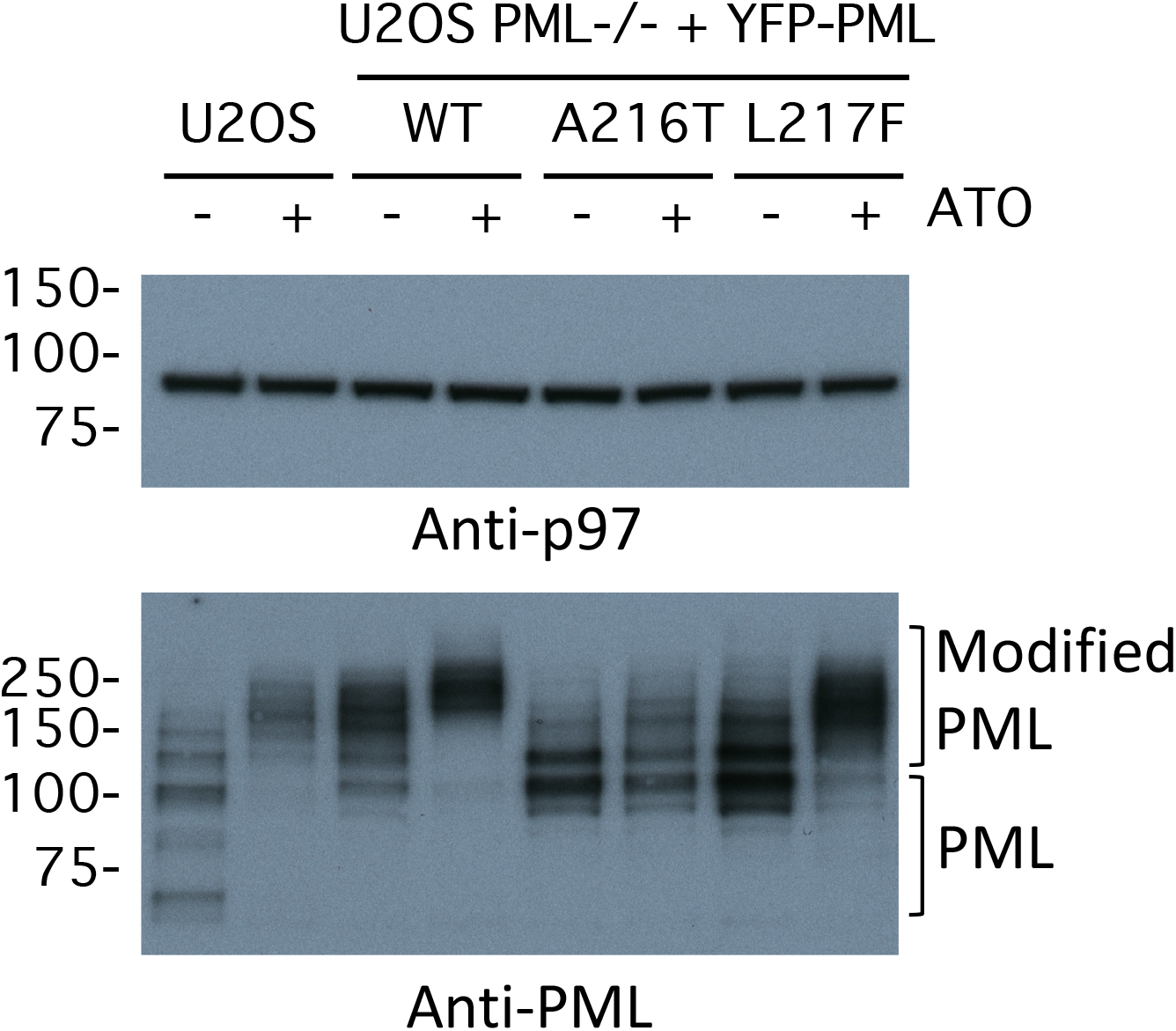
p97 expression levels do not differ between cell types or upon arsenic treatment. Western blots for p97 and PML of whole cell extracts from the indicated cell lines either treated or not with arsenic.

**Supplementary Figure 6.**
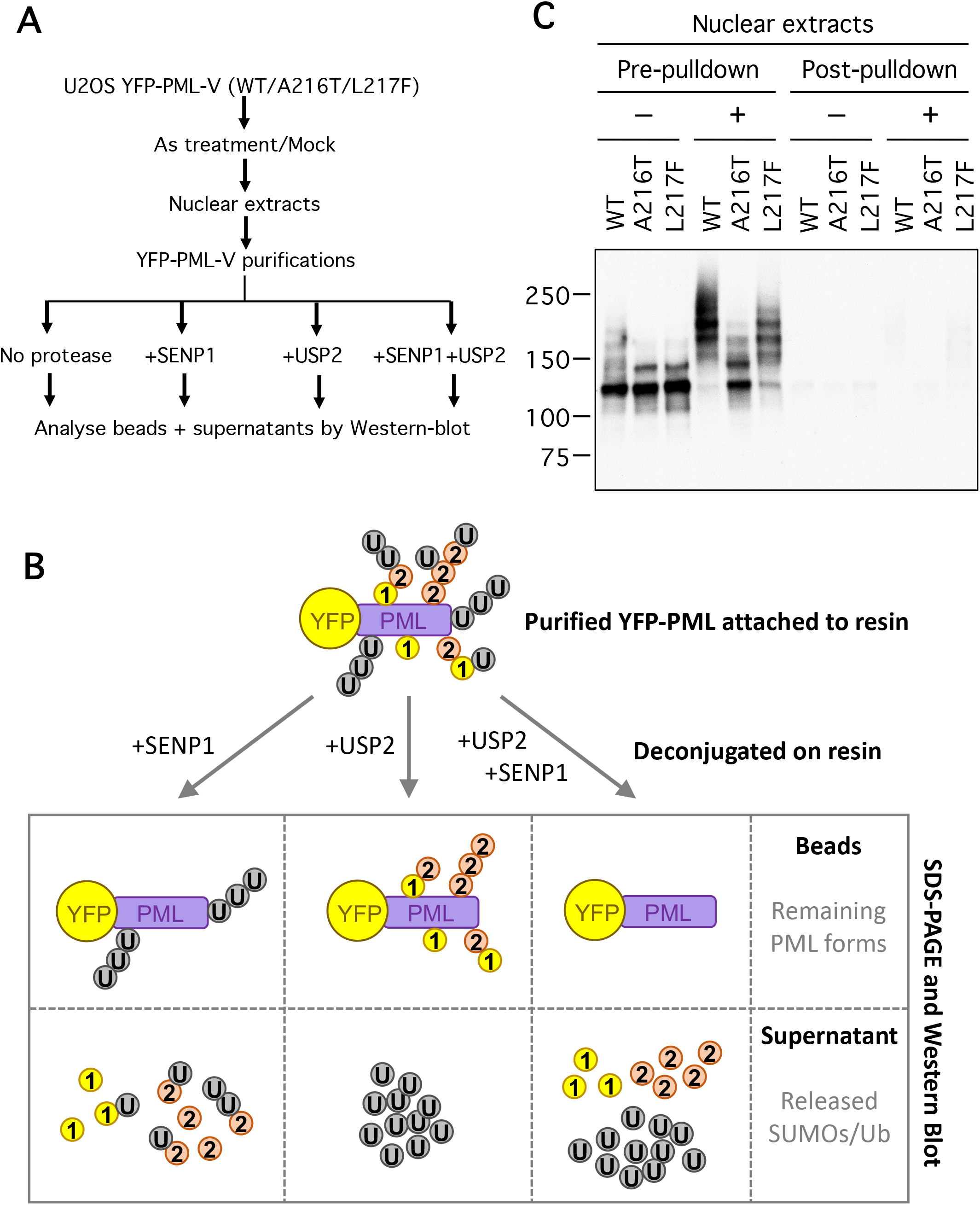
Treatment of purified PML bodies with recombinant SUMO and Ubiquitin proteases reveals the nature of PML post-translational modifications. A. Overview of the experimental design for detection of ubiquitin and SUMO conjugated to PML. B. Anti-PML Western blot for nuclear extracts from the indicated cell lines before and after purification of YFP-PML with GFP nanobody beads. C. Schematic depiction of the contents of YFP-PML bound to beads and resulting supernatants after treatment with specific proteases.

**Supplementary Figure 7.**
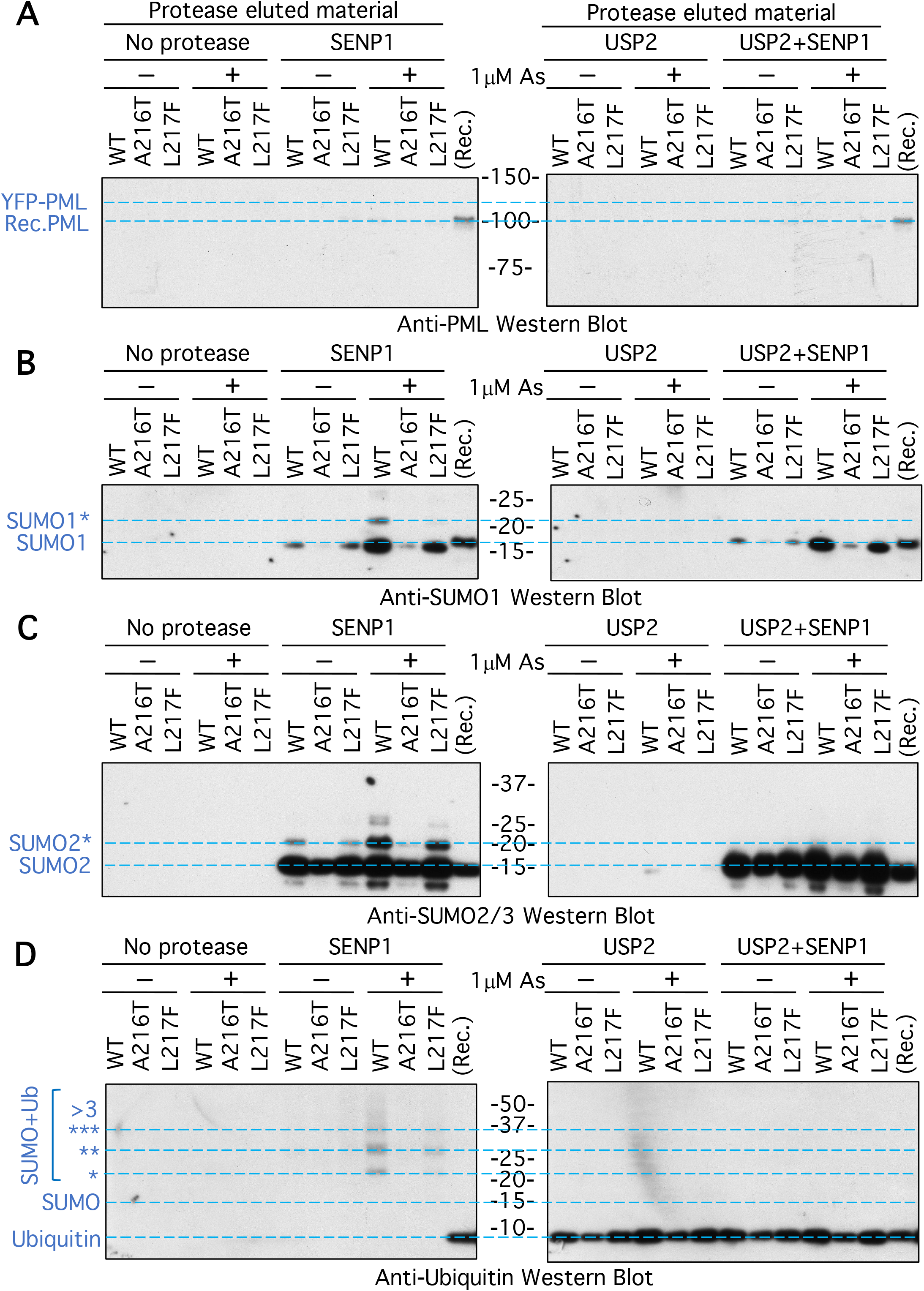
Western blot analysis of the material eluted by SENP1 and USP2 from the purified YFP-PML forms indicated. Associated with Figure 5. Western blots using antibodies specific for PML (A), SUMO1 (B), SUMO2/3 (C) and ubiquitin (D). 2ng standards of recombinant proteins (Rec.) were included with the gels. The positions of recombinant PML (Rec.PML), unmodified forms of YFP-PML, SUMO1, SUMO2 and ubiquitin are indicated, along with ubiquitinated species (*).

**Supplementary Figure 8.**
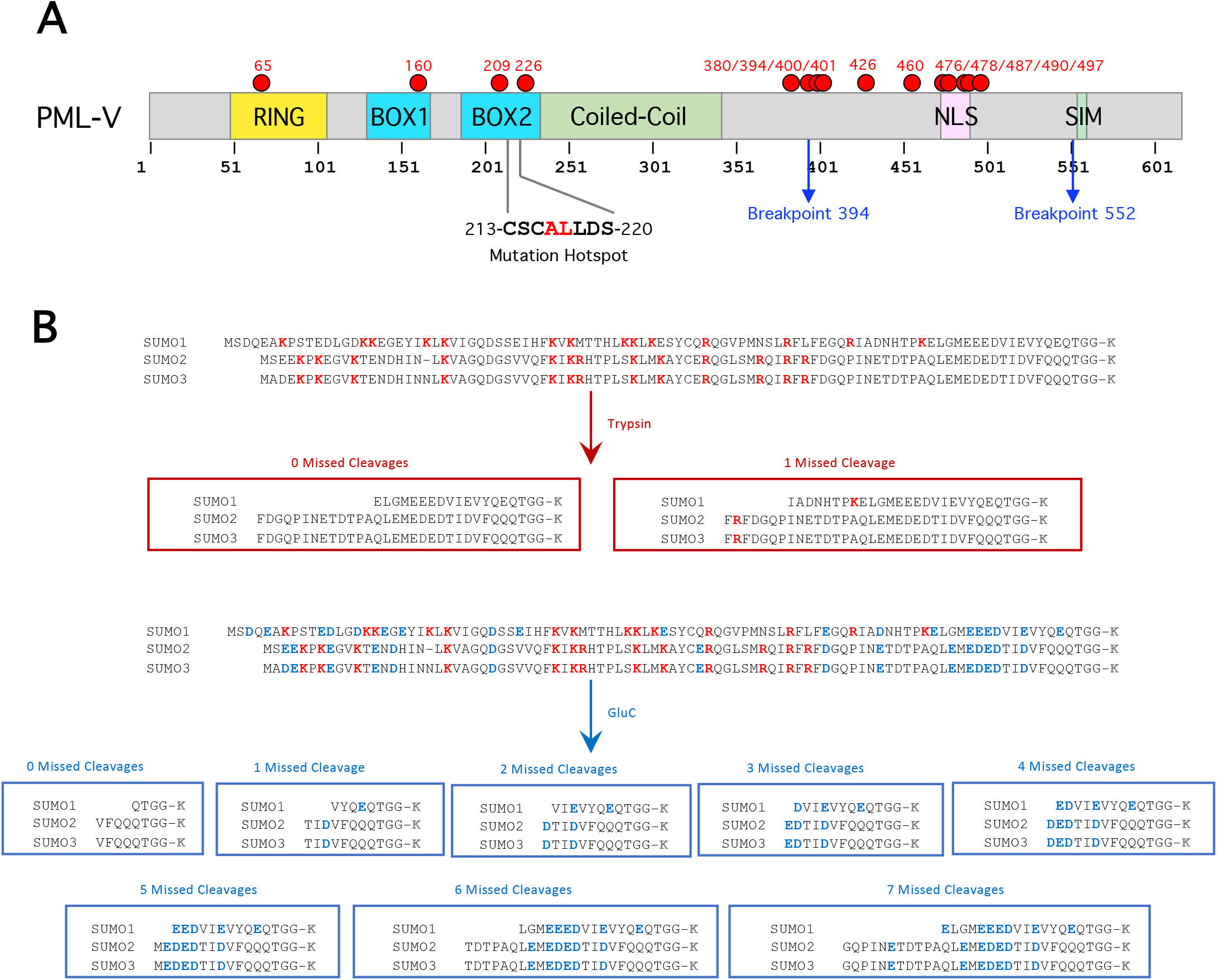
Known sites of PML SUMOylation and potential peptide adducts remaining on substrates after trypsin and GluC digestion. A. Positions of previously identified SUMO conjugation sites relative to the structural domains in PML. Isoform V is shown although identifications could be from other isoforms. Data taken from Hendriks et al 2017 (Hendriks, Lyon et al. 2017). Common breakpoints found in the PML-RARA and mutational hotspots found in arsenic insensitive forms are indicated. B. Overview of the potential remnants of SUMO C-termini attached to substrate lysine residues (K) when conjugates are cleaved by Trypsin, or GluC considering multiple missed cleavages.

**Supplementary Figure 9.**
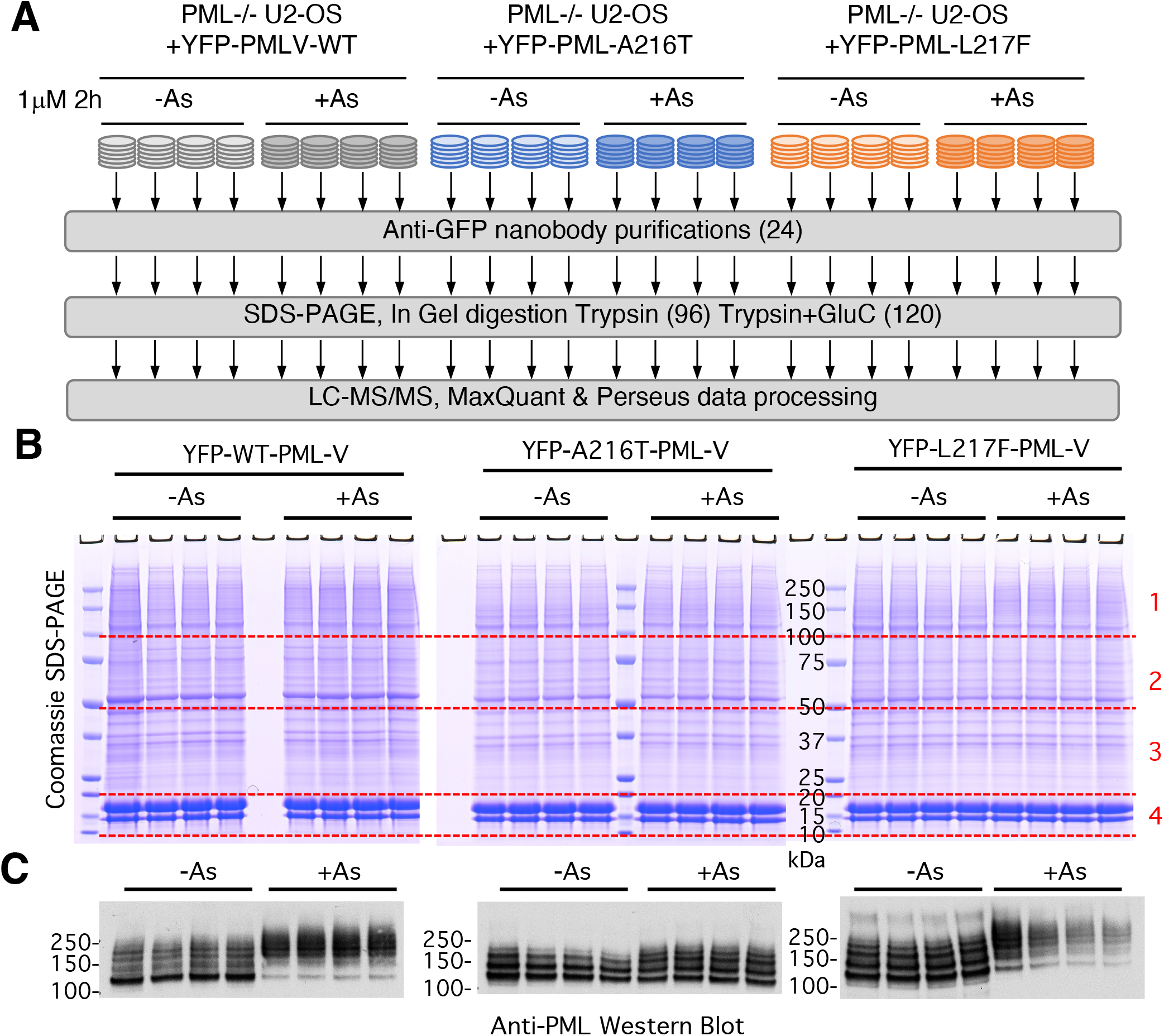
A proteomic study to quantitatively analyse arsenic-induced changes to PML post-translational modifications at the site level. A. Overview of the proteomic experimental design. B. Coomassie-stained gel showing the elutions from anti-GFP nanobody purifications for each replicate. Gel slices are numbered. C. Anti-PML western blot of a fraction of the anti-GFP nanobody purifications for each replicate.

**Supplementary Figure 10.**
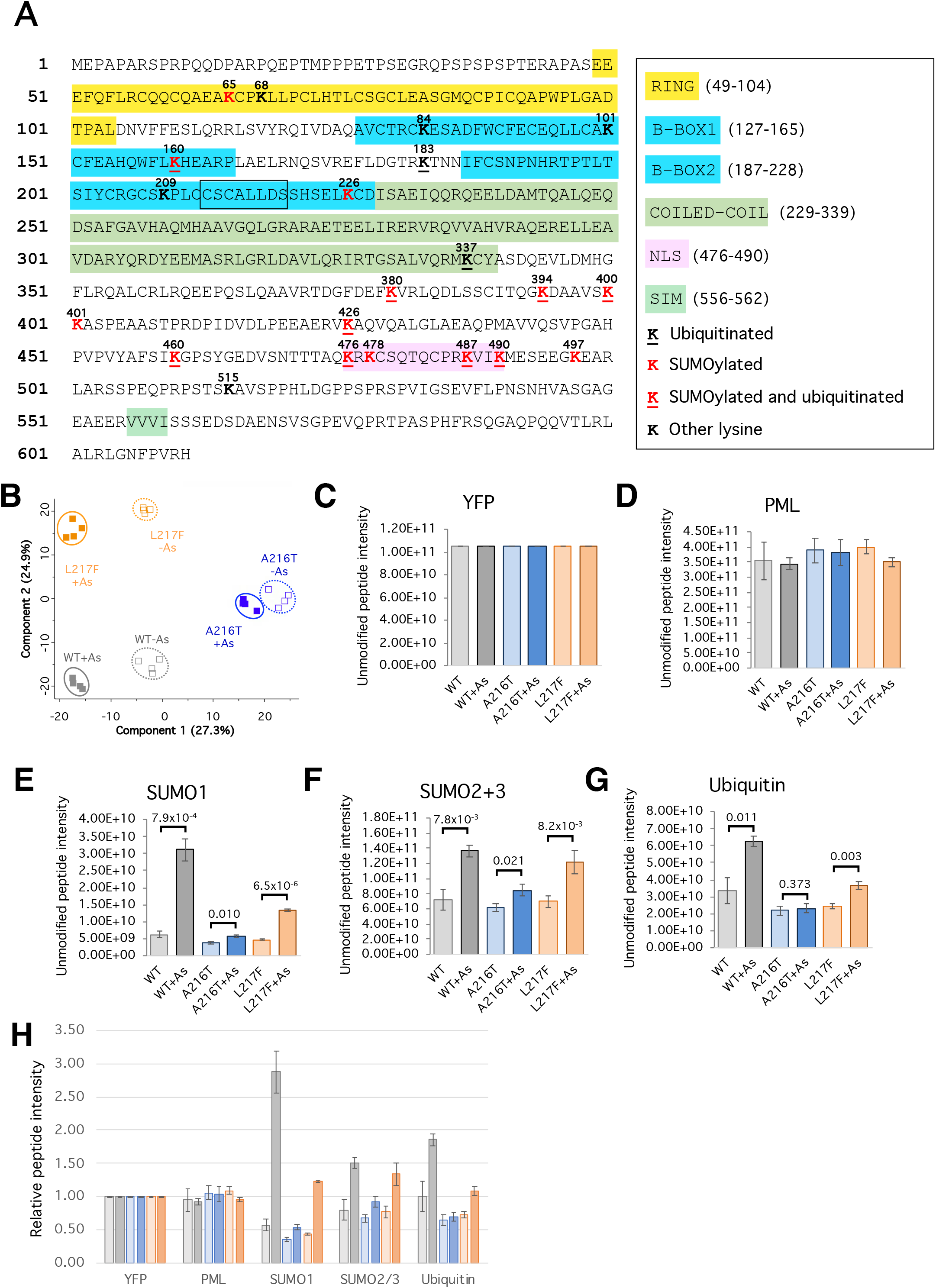
Overview of the proteomic data for the post- translational modification of PML WT, A216T and L217F. A. Summary of SUMO and ubiquitin conjugation sites identified in this study. Region of mutation hotspots is boxed. B. Principal component analysis of the quantitative proteomic data using only peptides from YFP-PML, SUMOs or ubiquitin. These included all unmodified peptides, phospho peptides, Ubiquitin branches and SUMO branches on YFP-PML, SUMO1, SUMO2/3 and Ubiquitin. Total is 343 peptides and uses log_2_ transformed and zero-replaced intensity values. C-G. Total unmodified peptide intensity data for the indicted proteins. Bars are average intensities with standard deviation error bars. p values are for two tailed student’s t-test comparing untreated with arsenic treated samples. n=4. H. Summary of the data shown in C-G normalized by the average intensity for each protein.

**Supplementary Figure 11.**
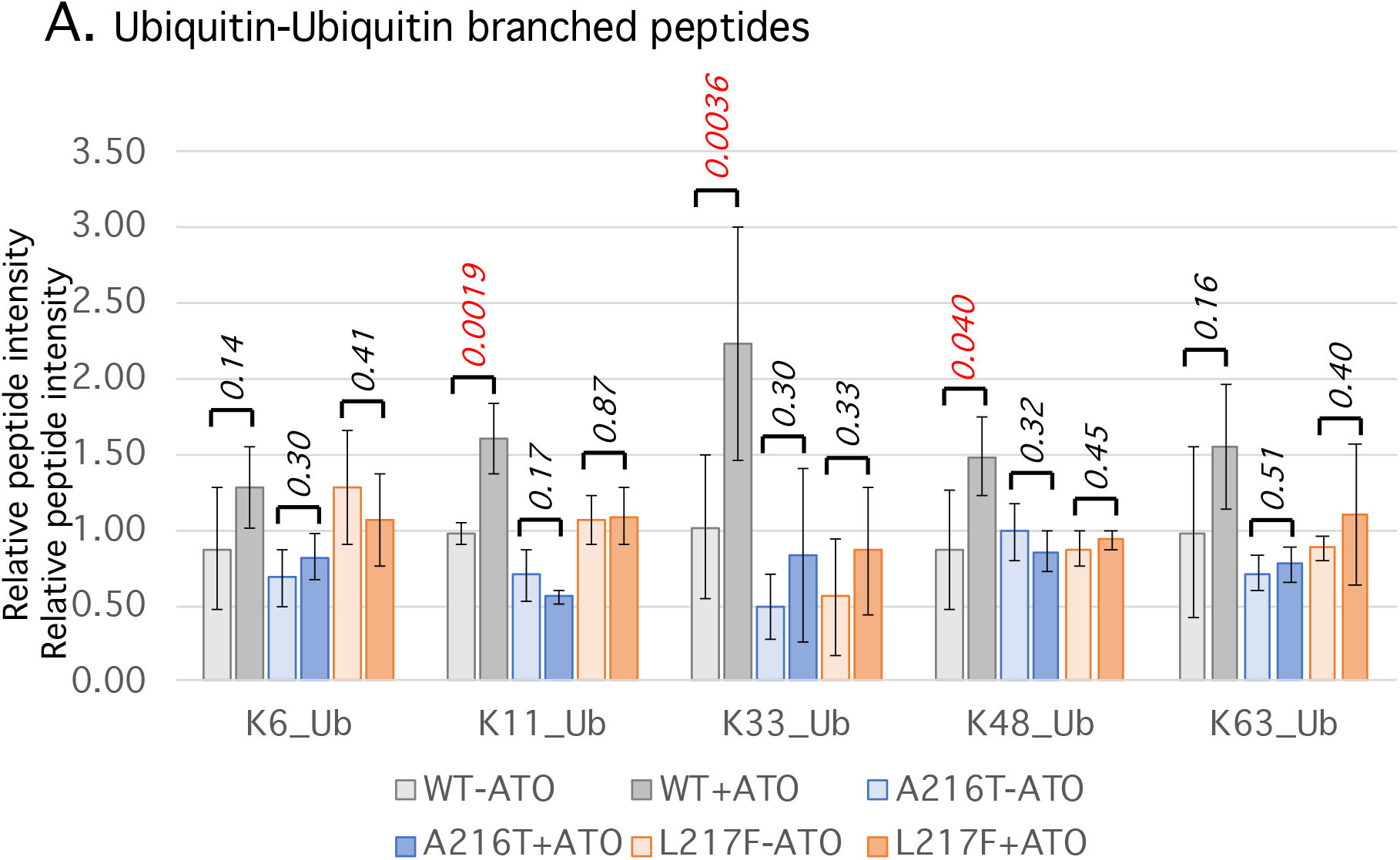
Column plots of relative peptide intensity data for selected peptide types. A. Quantitative data for peptides indicative of Ubiquitin- Ubiquitin linkages detected in the proteomics study. Columns are average values with SD error bars. Student’s t-test p-values are indicated.

**Supplementary Figure 12.**
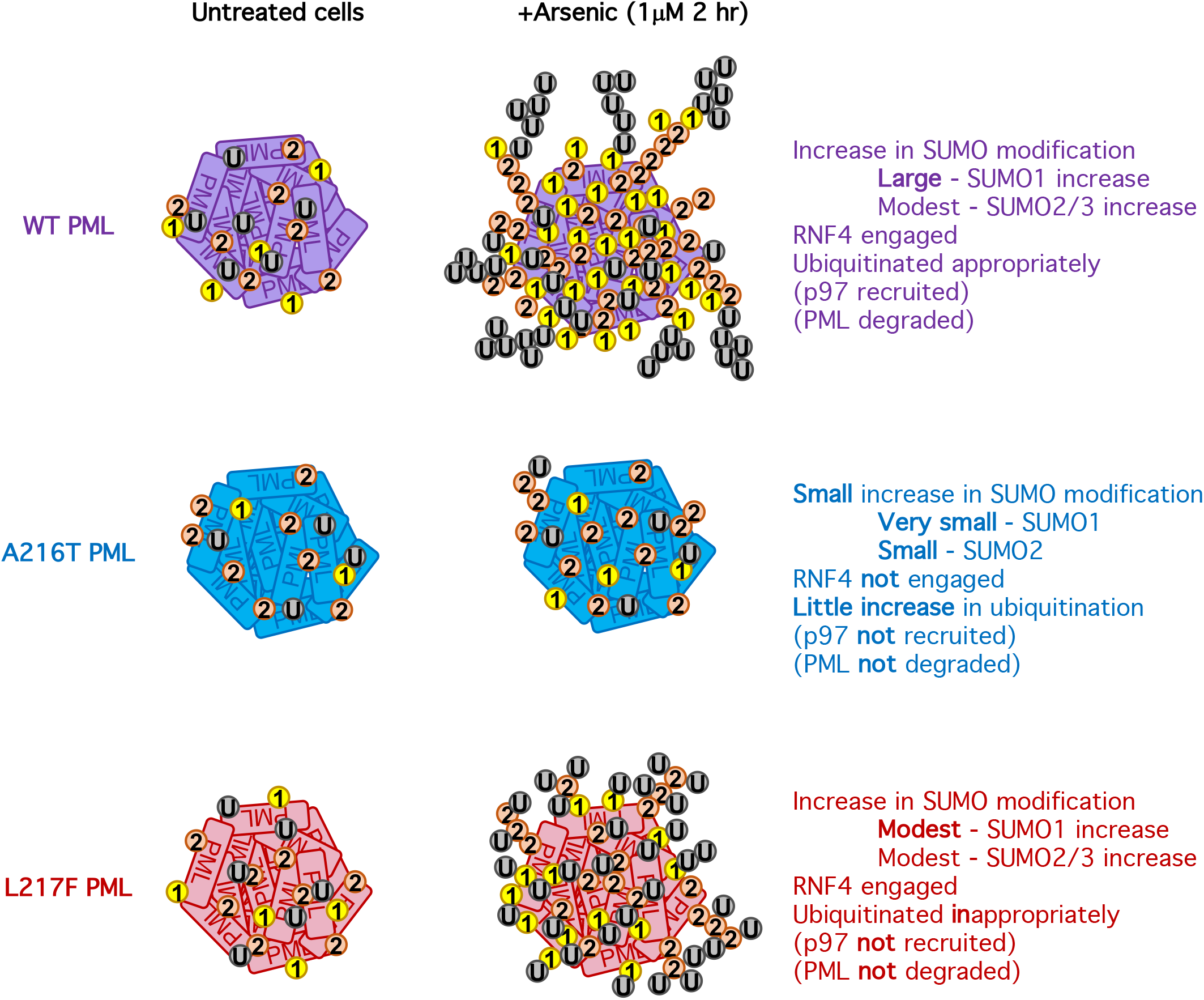
Schematic summary of the arsenic-induced responses of WT, A216T and L217F forms of PML. WT, A216T and L217F forms of PML are modified by broadly similar amounts of SUMO1 and SUMO2/3 in untreated cells. Exposure of cells to arsenic triggers an increase in SUMO2/3 conjugation for WT PML and L217F PML, and a much smaller increase for A216T PML. A large increase in SUMO1 conjugation is triggered for WT PML, with a smaller increase for L217F PML, and no significant SUMO1 change for A216T. Arsenic does not induce increased ubiquitination of A216T, while L217F experiences increased ubiquitination to PML and SUMO proteins. WT PML also becomes ubiquitinated on PML and SUMOs, but a fraction also forms ubiquitin-ubiquitin polymers. Neither A216T PML nor L217F PML develop the requisite SUMO and/or ubiquitin signals required to bind p97 and target them for degradation.

